# The CRL4B E3 ligase regulates mitosis by recruiting phospho-specific DCAFs

**DOI:** 10.1101/2022.10.14.512051

**Authors:** Anna Stier, Samuel Gilberto, Weaam I. Mohamed, Jonne Helenius, Ivan Mikicic, Tatjana Sajic, Petra Beli, Daniel J. Müller, Matthias Peter

**Affiliations:** Institute of Biochemistry, ETH Zurich, Otto-Stern-Weg 3, 8093 Zurich, Switzerland; Department of Biosystems Science and Engineering, ETH Zurich, Mattenstrasse 26, 4058 Basel, Switzerland; Institute of Molecular Biology, Ackermannweg 4, 55128 Mainz, Germany; Institute of Molecular Systems Biology, ETH Zürich, Otto-Stern-Weg 3, 8093 Zürich, Switzerland; Institute of Developmental Biology and Neurobiology (IDN), Biozentrum 1, Hanns-Dieter-Hüsch-Weg 15, Johannes Gutenberg University, 55128 Mainz, Germany

## Abstract

The cullin-4 paralogs CUL4A and CUL4B assemble E3 ubiquitin ligase complexes regulating multiple chromatin-associated cellular functions. Although they are structurally similar, we found that the unique N-terminal extension of CUL4B is heavily phosphorylated during mitosis, and the phosphorylation pattern is perturbed in the CUL4B-P50L mutation causing X-linked intellectual disability (XLID). Phenotypic characterization and mutational analysis revealed that CUL4B phosphorylation is required for efficient progression through mitosis, controlling spindle positioning and cortical tension. Interestingly, while CUL4B phosphorylation triggers chromatin exclusion, it critically promotes binding to actin regulators and two previously unrecognized, CUL4B-specific DCAFs, LIS1 and WDR1. Indeed, co-immunoprecipitation experiments and biochemical analysis revealed that LIS1 and WDR1 interact with DDB1, but their binding requires the phosphorylated N-terminal domain of CUL4B. Together, our study uncovers previously unrecognized DCAFs relevant for mitosis and brain development that specifically bind CUL4B, but not the CUL4B-P50L patient mutant, by a phosphorylation-dependent mechanism.

## Introduction

Cullin-RING E3 ligases (CRLs) comprise the largest family of RING-based E3 ubiquitin ligases and catalyze approximately 20% of all ubiquitination reactions leading to proteasomal degradation in mammalian cells^1^. CRLs use one of eight cullins (CRL1-3, CRL4A-B, CRL5, CRL7, CRL9) as a scaffolding subunit for their modular assembly. CRL activity depends on the ubiquitin-like modifier NEDD8 and additional mechanisms altering substrate recruitment, highlighting the high complexity of CRL regulation^2–5^.

CUL4-based E3 ligases regulate many cellular activities, including chromatin-associated processes like DNA damage repair^6–9^ and cell cycle progression^10–13^. CRL4 complexes engage DDB1-CUL4 associated factors (DCAFs) as substrate receptors (SRs), which are characterized by WD40 domains and a helix-loop-helix motif mediating their binding to the adaptor DDB1^9, 14^. Interestingly, vertebrates encode two conserved paralogs, CUL4A and CUL4B, with *CUL4B* gene located on the X-chromosome. The paralogs share over 80% sequence identity and all functional domains fold into nearly identical structures.

The major difference between CUL4A and CUL4B is an extended N-terminus unique to CUL4B, which is mostly unstructured and thus commonly truncated in biochemical studies^9, 15^. As both cullins are widely expressed and use DDB1 to bind a common set of SRs, CRL4A and CRL4B complexes are thought to function in a redundant manner^6, 16, 17^. Nevertheless, several studies report CRL4A or CRL4B-specific functions in diverse cellular contexts^18^. Indeed, while targeted MS quantification confirmed that CUL4A and CUL4B equally interact with most previously identified DCAFs, a small subset was found to preferentially bind CRL4A or CRL4B complexes. For example, CRBN prefers to assemble with CRL4A, while AMBRA1 favors CRL4B^19^. However, the underlying reason for this DCAF preference is not understood.

Distinct functions of CRL4A and CRL4B complexes are also apparent from studies in mice. CUL4B is essential during mouse embryogenesis, since deletion leads to cell cycle defects and apoptosis in extra embryonic tissues^20, 21^. CUL4A knockout mice, however, show no effects on embryonic viability but fail spermatogenesis^22, 23^. Moreover, CUL4B knockouts in mouse epiblasts identified specific neuronal and behavioral defects^24^. In line with these findings, mutations in human CUL4B but not CUL4A are associated with X-linked intellectual disability (XLID)^25–28^, consistent with non-redundant functions of the two cullin scaffolds in brain development^29^. While most patient mutations result in loss of CRL4B ligase activity, a single point mutation of CUL4B (P50L) mapping to its unique N-terminus also causes XLID^28^, but the underlying molecular mechanism remains unclear.

Here, we examined the differential roles of CUL4A and CUL4B in cell cycle progression. We found that cells lacking CUL4B show mitotic spindle positioning defects and reduced cortical tension. Molecular analysis revealed that CUL4B is specifically phosphorylated during mitosis, which increases its interaction with many actin regulators and two previously unrecognized DCAFs, LIS1 and WDR1. Indeed, cells expressing phosphorylation-defective CUL4B mutants including the patient-derived XLID mutant P50L exhibit mitotic defects and the mutant proteins fail to interact with LIS1 and WDR1. Importantly, *in vitro* reconstitution experiments confirm that LIS1 and WDR1 bind DDB1, but also require a second, phosphorylation-dependent interface with the unique amino terminal domain of CUL4B. Our study thus provides a molecular mechanism explaining CRL4B specificity with implications for brain development and identifies a previously unrecognized subset of DCAFs that bind CUL4B in a phosphorylation-dependent manner during mitosis.

## RESULTS

### CRL4B-specific functions in mitosis

To investigate functional differences between CRL4A and CRL4B (Fig. 1A), we used RNAi to knock down either CUL4A, CUL4B or DDB1 in HeLa Kyoto cells, with specific downregulation verified by immunoblotting (Fig. 1B). Upon RNAi-depletion of DDB1, CUL4A and CUL4B accumulated predominantly in their deneddylated, inactive form. In contrast, neddylated CUL4A accumulated upon depletion of CUL4B, and conversely neddylated CUL4B increased with CUL4A depletion, implying a potential compensation mechanism (Fig. 1C). Despite this paralog compensation, MTT assays revealed decreased growth rates of approximately 30% for CUL4A and 20% for CUL4B RNAi, respectively, compared to control cells (Fig. 1D). The growth rate of CRISPR/Cas9 CUL4B knockout cells (ΔCUL4B) was similarly reduced (Extended Data Fig. 1A, B, C). This growth delay phenotype was enhanced to 40-50% when combining CUL4A and CUL4B depletion or upon the depletion of the common adaptor DDB1, implying redundant as well as non-redundant functions of CRL4A and CRL4B for cell growth.

**Figure 1:**
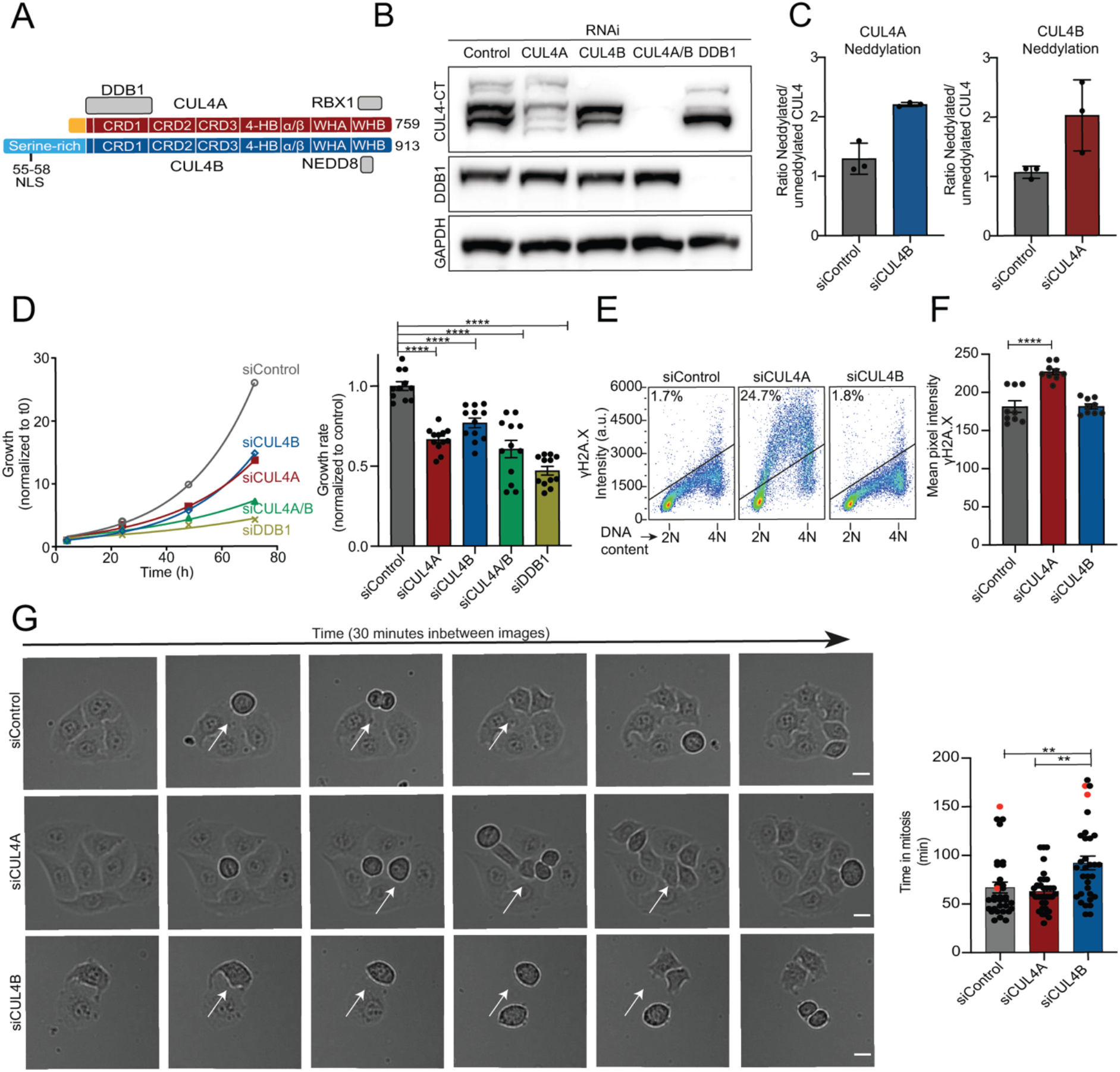
CUL4B-specific functions in mitosis. (**A**) Schematic representation of structural and functional domains of CUL4A (top, red) and CUL4B (bottom, blue) following the annotation described in Fischer *et al*.^9^. The common CRL4 subunits and their binding sites are indicated in grey. (**B**) Western blot of RNAi efficiency and specificity for oligos used in (C-G). HeLa Kyoto cells were incubated for 72 h with control oligos or RNAi-oligos depleting CUL4A, CUL4B or DDB1. Specific downregulation of the indicated proteins was analyzed by immunoblotting. GAPDH controls for equal loading. (**C**) Quantification western blot bands and the ratio between neddylated vs unneddylated CUL4 in absence of the paralog is calculated. Graphs displays mean with SEM of N=3 (**D**) Growth curves and growth rates of RNAi depleted HeLa Kyoto cells measured by MTT assay. Graph represents mean with SEM after 72 h normalized to siControl N=3 x 3 technical replicates. (**E**) Flow cytometry analysis of Hela cells treated with the indicated RNAi for 72h and immune-stained for γH2A.X, DNA staining with propidium iodide, N=3. (**F**) DNA damage levels analyzed by immunofluorescence staining of γH2A.X in RNAi depleted HeLa Kyoto cells. Quantification of mean pixel intensity of γH2A.X signal in the nucleus. Graph displays mean with SEM of N= 3 x 3 technical replicates of at least 50 cells in each experiment. (**G**) Live cell imaging of mitosis in RNAi depleted HeLa Kyoto cells. Brightfield images were taken every 3 min, and mitotic duration was quantified by measuring the time from cell rounding to telophase. Mean with SEM is depicted for N=3 experiments analyzing at least 10 cells. Red symbols mark incomplete mitotic events during the movie. Statistical analysis for all graphs was performed with 1-ANOVA and Dunnett’s multiple comparison. White arrows mark cells undergoing mitosis. Scale bar: 10 μm. See also Extended Data Fig 1.

Since CRL4 E3 ligase activity is important in DNA replication and DNA damage repair^7^, we hypothesized that the growth delay might be caused by accumulation of DNA damage. Indeed, the number of cells positive for phosphorylated histone H2A.X (γH2A.X) increased more than 10-fold following CUL4A depletion (Fig. 1E). Similarly, γH2A.X staining by immunofluorescence showed an increase of approximately 25% in cells RNAi-depleted for CUL4A (Fig. 1F, Extended Data Fig. 1E). In contrast, γH2.AX levels in CUL4B-depleted or ΔCUL4B cells were unchanged relative to wild-type controls (Fig. 1E, F, Extended Data Fig. 1D, F), implying that the observed growth delay is caused by a distinct CRL4B function.

We next used brightfield live-cell microscopy to visualize division of cells lacking CUL4A or CUL4B. Interestingly, cells lacking CUL4B exhibit a significant lengthening of mitotic duration, which was not observed in CUL4A-depleted or control cells (Fig. 1G, Extended Data Fig. 1G). Indeed, cells with impaired CRL4B took on average 15-25 minutes longer compared to control cells when measured from rounding-up upon mitotic entry until the onset of cytokinesis. Consequently, we hypothesized a CRL4B-specific function during mitosis as a possible cause for the observed growth delay of CUL4B depleted cells. Taken together, these results suggest that CRL4B has a unique function that is important for efficient execution of mitosis, explaining the observed growth delay of CUL4B depleted cells.

### The P50L patient mutation alters mitotic-specific phosphorylation of CUL4B’s unique N-terminal extension

We examined CUL4B expression and localization through the cell cycle. As expected, CUL4A and CUL4B are expressed throughout the cell cycle, and their neddylated, active forms are present at all stages. However, when cells were arrested in mitosis either by taxol or nocodazole, CUL4B migrated significantly slower on SDS-PAGE (Fig. 2A), indicative of a post-translational modification. Indeed, slower-migrating CUL4B species were readily reversed by incubating mitotic extracts with lambda-phosphatase (λ-PP) suggesting phosphorylation (Fig. 2B). No mitotic phosphorylation could be detected for CUL4A or when CUL4B was analyzed in extracts prepared from cells arrested in S-phase with thymidine or G2 with RO-3306. CUL4B phosphorylation was also monitored after a mitotic shake-off in asynchronized cells (Fig. 2C) or in cells released from a G2 block, thus synchronously progressing from G2 into mitosis (Extended Data Fig. 2A). In both cases, slower-migrating CUL4B was observed, demonstrating that CUL4B but not CUL4A is specifically phosphorylated during mitosis. Moreover, phosphorylation sites during mitosis were contained in the unique N-terminus of CUL4B, since deletion of the unique N-terminus of CUL4B abolished the phosphorylation of CUL4B (Extended Data Fig. 2B).

**Figure 2:**
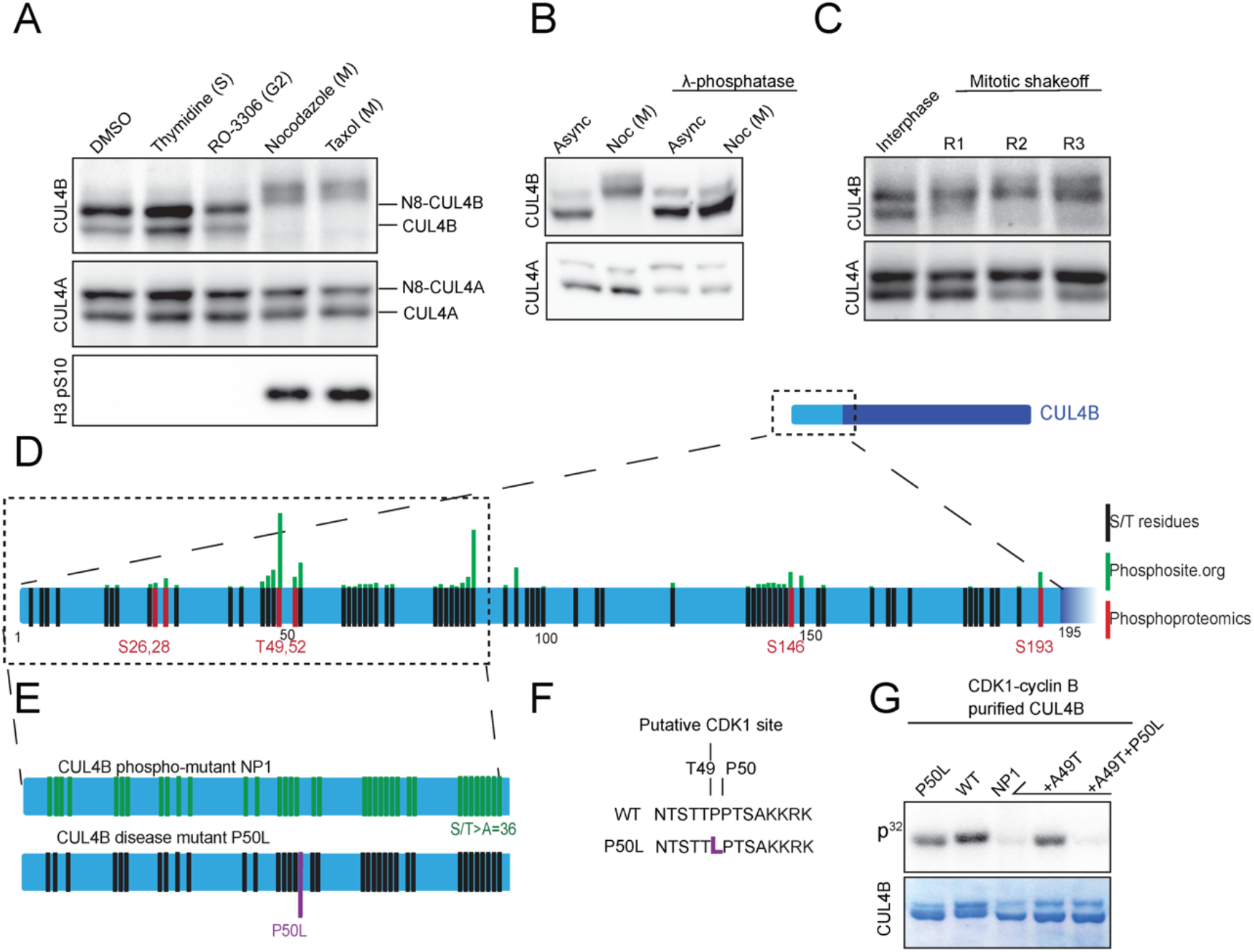
N-terminal phosphorylation of CUL4B during mitosis. (**A**) CUL4A and CUL4B were analyzed by immunoblotting of extracts prepared from HeLa Kyoto cells arrested in S-phase (S= thymidine), G2 (R0-3306) or mitosis (M= nocodazole or taxol) (N=3). H3 pS10 (phosphorylated histone H3 at serine 10) was used as a mitotic marker. (**B**) CUL4A and CUL4B were immunoblotted of extracts prepared from asynchronous (Asyn) or nocodazole (Noc)-arrested mitotic cells and treated as indicated with λ-phosphatase (N=3). (**C**) Mitotic population of unperturbed cells was harvested by shake-off. Three subsequent harvests are depicted (R1-R3). The interphase population are cells that remained attached to the culture dish. (**D**) Annotation of the serine and threonine residues on the N-terminus of CUL4B (black). Green lines represent phospho-sites found on Phosphosite.org; the length indicates the number of studies identifying this site. The red sites were identified by phosphoproteomics on immunoprecipitated CUL4B from nocodazole-arrested mitotic cells (N=2). All sites reported by phospho-proteomics have a localization probability >0.75 except T52 (0.68). (**E**) Representation of CUL4B amino-terminal mutants used in this study. In the NP1 mutant the green serine and threonine residues located in the first 90 amino acids were mutated to non-phosphorylatable alanine. The P50L XLID patient mutation is highlighted in purple ^28^. (**F**) Representation of the putative T49 CDK1 consensus site in WT CUL4B, which is altered in the P50L mutant. (**G**) *In vitro* kinase assays with purified CDK1-cyclinB and Sf9-purified CUL4B and mutant proteins in the presence of [γ-^32^P]ATP. The autoradiogram (top panel) shows CUL4B phosphorylation, while coomassie staining (lower panel) controls equal loading of CUL4B and mutants (N=3 for WT, P50L; NP1; N=2 for NP1-A49T and NP1-A49T-P50L). Note that the A49T mutation in the NP1 mutant restores phosphorylation of the T49 residue, but only in the presence of the adjacent proline at position 50. See also Extended Data Fig. 2.

Searching the phosphosite.org database revealed many phospho-sites located in the unique amino-terminal extension of CUL4B (Fig. 2D). To systematically identify the mitotic phosphorylation sites, we immunoprecipitated CUL4B from mitotic cells and mapped phosphorylated peptides after TiO_2_-enrichment using mass-spectrometry. To facilitate efficient immunoprecipitation without overexpression, we replaced endogenous CUL4B with a functional HA-tagged CUL4B fusion (Extended Data Fig. 2 C, D). Phospho-site enrichment and quantitative mass spectrometry identified several sites (S26,28; T49,52; S193) in the unique N-terminus of CUL4B, which are phosphorylated during mitosis (Fig. 2D, Table S1). Importantly, T49 emerged as a major mitotic site, which conforms to the minimal consensus site for proline-directed kinases such as cyclin-dependent kinases (CDKs) (Fig. 2E, F). Indeed, *in vitro* kinase assays in the presence of [γ-^32^P]ATP (Extended Data Fig. 2E) demonstrated that purified CUL4B, but not CUL4A, was readily phosphorylated by CDK1-cyclin B, and to a lesser extent also by PLK1 and CK1.

Since T49 is located next to the P50L mutation found in XLID patients, we performed CDK1 *in vitro* kinase assays using purified CUL4B-P50L as a substrate. In addition, we analyzed a CUL4B mutant (NP1) that has all serine and threonine sites within the first ninety amino acids of the N-terminus mutated to non-phosphorylatable alanine residues (Fig. 2E). Indeed, CDK1-mediated phosphorylation of the P50L mutant was reduced compared to wild-type (WT), and almost abolished with the NP1 mutant (Fig. 2G). Importantly, re-addition of the T49 phospho-site to the NP1 mutant partially restored a P^32^ signal, while T49 phosphorylation was strongly reduced when combined with the P50L mutation. We conclude that the unique amino terminal extension of CUL4B is phosphorylated on multiple sites during mitosis and that CDK1-cyclin B is the main CUL4B kinase *in vitro*. A major mitotic phospho-site is T49, and its phosphorylation is strongly diminished by the P50L patient mutation.

### Mitotic CUL4B phosphorylation promotes progression through mitosis and regulates chromatin association

In CUL4B knockout HeLa Flp-in cells (ΔCUL4B_Flp_), we quantified mitotic duration when complemented with inducible expression of CUL4B WT, or the phosphorylation-defective mutants P50L or NP1 by live-cell microscopy. The NP1 and P50L mutant proteins were expressed at comparable levels, while expression of CUL4B WT was slightly lower (Fig. 3A). The mutations neither altered expression levels nor the neddylation status of CUL4A or CUL4B. However, expression of both phosphorylation-defective mutants NP1 and P50L failed to rescue the mitotic delay of cells lacking CUL4B, and mitotic duration remained around 90 minutes in contrast to 60 minutes observed for CUL4B WT controls (Fig. 3B, Extended Data Fig. 3A). Together, these results demonstrate that mitotic CUL4B phosphorylation and in particular the T49 site is important for mitotic function.

**Figure 3:**
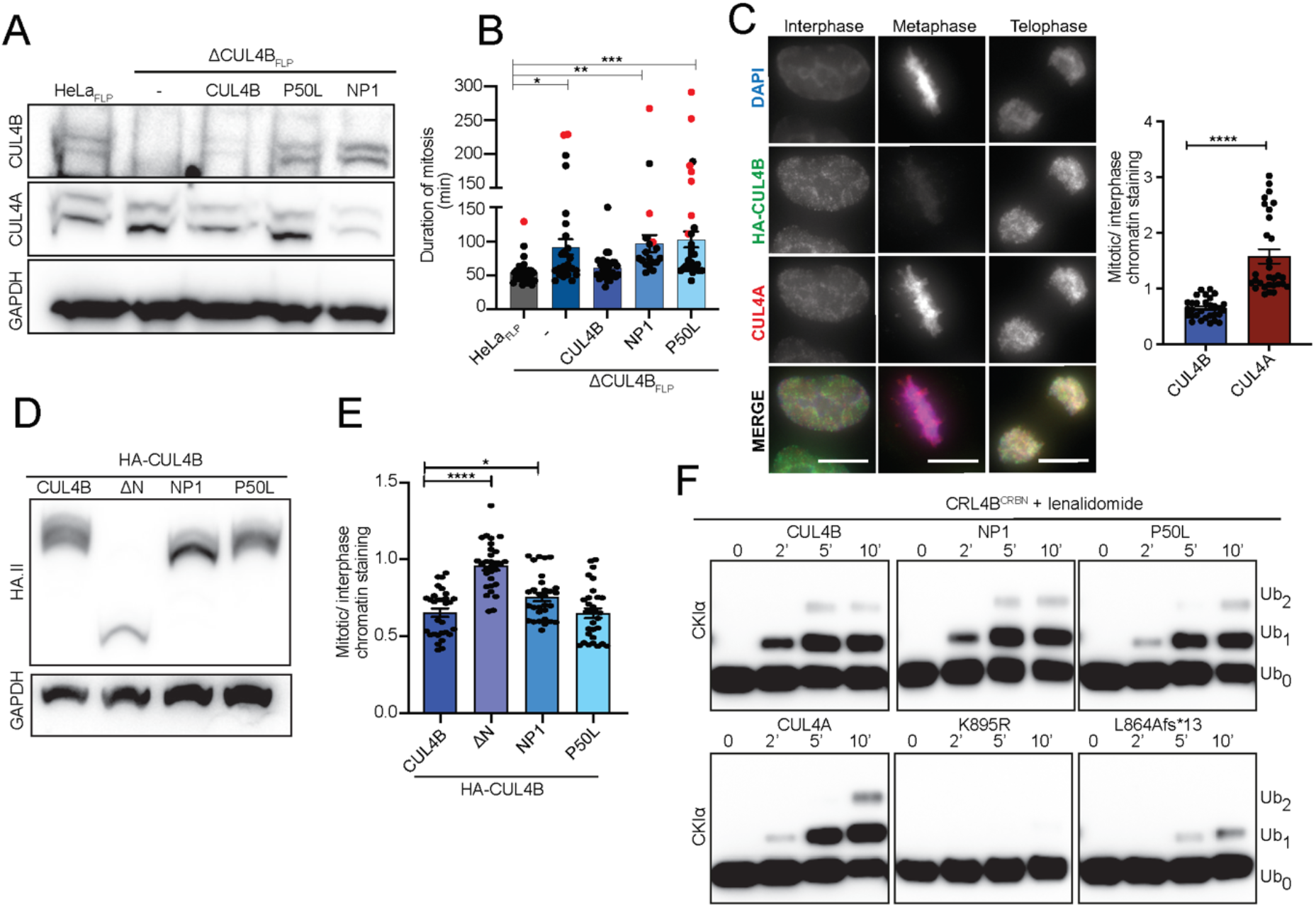
Mitotic CUL4B phosphorylation promotes progression through mitosis and regulates chromatin association. (**A**) Western blot analysis to monitor CUL4A or CUL4B levels in extracts prepared from HeLaFLP control cells or HeLaFLP cells deleted for CUL4B by CRISPR/Cas9 (ΔCUL4B_FLP_) but stably expressing no CUL4B (-), CUL4B or the indicated NP1 or P50L mutants from the doxycycline-inducible promoter. GAPDH was included to control equal loading. (**B**) Live cell imaging in the brightfield channel from cell rounding to telophase was used to quantify the mitotic duration of HeLaFLP control cells or ΔCUL4B_FLIP_ cells expressing CUL4B, NP1 or the P50L mutant after doxycycline induction. Mean with SEM are depicted of N= 3 independent experiments with at least 10 cells. Red symbols mark cells with incomplete mitosis during the movie. (**C**) Immunofluorescence analysis of CUL4A and HA-CUL4B in interphase, metaphase, or telophase cells after cytoplasmic pre-extraction. The DNA was visualized by DAPI. Representative images show maximum intensity Z-projections of the acquired Z-stacks. The contrast settings are identical between images of a single channel. Scale bar: 10 μm. The mean grey values of HA-CUL4B or CUL4A on metaphase plates were measured and normalized with ImageJ to mean grey values of the nuclear signal in interphase nuclei. The graph represents the mean with SEM for normalized metaphase cells in N= 3 independent experiments with at least 10 cells. (**D**) The expression levels of endogenous HA-tagged WT and the indicated CUL4B mutants were compared by immunoblotting with HA.II antibody. GAPDH controls for equal loading (**E**) Chromatin exclusion of HA-tagged WT and the indicated HA-CUL4B mutants in metaphase was quantified by immunofluorescence after cytoplasmic pre-extraction and normalized with ImageJ to mean grey values of the nuclear signal in interphase cells. The graph represents the mean with SEM for the normalized metaphase signal from N= 3 independent experiments with at least 10 metaphase plates. (**F**) *In vitro* E3 ligase activity assays using the indicated reconstituted CRL4B^CRBN^ complexes and the neo-substrate CKlα. At time 0, lenalidomide was added to start the reaction and CKlα ubiquitination was followed at the times indicates (in minutes) by immunoblotting with CKlα antibody. N≥3 for CUL4B, CUL4A, P50L, K895R and L864fs*13; N=1 for NP1. Statistical analysis was performed with 1-ANOVA and Dunnett’s multiple comparison or t-test. Note that the *in vitro* E3 ligase activity of CRL4B and the P50L patient mutation is comparable. See also Extended Data Fig. 3.

To test the underlying mechanism, we used immunofluorescence to compare chromatin recruitment of endogenous CUL4A and CUL4B during the cell cycle. To simultaneously observe the two CUL4 variants, we used the endogenously HA-tagged CUL4B cell line and co-stained with antibodies specific for CUL4A and the HA-tag. We also employed pre-extraction to decrease cytoplasmic CUL4 signal. While both CUL4A and HA-CUL4B are present in the nucleus during interphase, we observed a 2-fold reduction of CUL4B signal on mitotic chromatin, while the CUL4A signal slightly increased at the chromatin in metaphase cells (Fig. 3C). Cellular fractionation assays of asynchronous or mitotic cells consistently showed that CUL4B was excluded from mitotic chromatin, while CUL4A remained bound to chromatin throughout mitosis. Chromatin recruitment for CUL4A and CUL4B was abolished after RNAi of DDB1 (Extended Data Fig. 3B), consistent with earlier reports that CRL4 chromatin recruitment is mediated via specific substrate receptors like DDB2 or RepID^30, 31^.

To test whether exclusion of CUL4B from mitotic chromatin is due to phosphorylation, we used endogenously HA-tagged CUL4B mutant cell lines (Fig. 3D). Immunofluorescent staining after cytoplasmic pre-extraction revealed that a N-terminal truncated mutant of CUL4B (ΔN) remains bound to chromatin, comparable to CUL4A localization. Likewise, the NP1 mutant is partially retained, but no chromatin dissociation defect could be detected for the P50L mutant (Fig. 3E). Cellular fractionation assays of mitotic cells transiently expressing Flag-tagged CUL4B, or the two phospho-mutants confirmed slight retention of the NP1 mutant, while the P50L mutant dissociated from mitotic chromosomes comparable to CUL4B WT (Extended Data Fig. 3C). We conclude that while phosphorylation of the unique amino-terminal domain triggers dissociation of CUL4B from mitotic chromosomes, this effect cannot solely explain the function of CUL4B in regulating mitosis.

Since most CUL4B mutations found in XLID patients decrease CRL4B activity or protein stability, we next tested whether phosphorylation of the amino-terminal domain regulates ubiquitination activity. We assembled recombinant CRL4 complexes with CUL4A, CUL4B or the phosphorylation-defective P50L or NP1 mutants. Additionally, we included the neddylation-defective CUL4B-K895R and the L864Afs*13 disease mutant. We monitored lenalidomide-induced CKIα ubiquitination to quantify CRL4B^CRBN^ activity *in vitro*. As shown in Figure 3F, no significant difference in CKIα ubiquitination kinetics could be detected between CRL4B WT, the phosphorylation-defective CRL4B mutants and CRL4A, implying that phosphorylation of the amino-terminal domain is not required for E3 ligase activity. In contrast, the L864Afs*13 disease mutant strongly decreased CKIα ubiquitination, while ubiquitination activity of the CUL4B-K895R mutant was completely abolished. Taken together, these data indicate that after release from mitotic chromosomes, phosphorylated CUL4B may associate with novel factors to regulate mitotic progression.

### LIS1 and WDR1 function as unconventional, phospho-dependent CRL4B DCAFs during mitosis

To identify proteins interacting with CUL4B in a phosphorylation-dependent manner during mitosis, we set up a SILAC-based mass spectrometric approach (Fig. 4A). Briefly, HA-tagged CUL4B or the phosphorylation-defective P50L or NP1 mutants were immunoprecipitated from SILAC-labeled cells arrested in mitosis with nocodazole. DCAF exchange during lysis was inhibited by blocking the NEDD8 cycle by addition of the CSN-(CSN5i-3) and NAE inhibitors (MLN4924)^19^. Lysates were immunoprecipitated with HA.II-antibodies and mixed before the last wash. Bound proteins were separated by SDS-PAGE and identified by LC-MS/MS. As expected, this approach verified that essential components like NEDD8, DDB1 and RBX1 as well as the SR exchange factor CAND1 bind CUL4B by a phosphorylation-independent mechanism. Likewise, we identified CUL4A as a strong interactor of CUL4B and phosphorylation-defective mutants, indicating the formation of heterodimers or oligomeric assemblies of CRL4A and CRL4B (Fig. 4B, Table S2). Importantly, however, this analysis also revealed several proteins particularly enriched in the phospho-specific interactome of CUL4B. Most notably, we discovered LIS1 and WDR1, which interact with WT CUL4B but show reduced binding to phosphorylation-defective P50L or NP1 mutants. Both proteins possess WD40 domains followed by a putative helix-loop-helix motif characteristic for established DCAFs (Fig. 4B, Extended Data Fig. 4A)^9, 32^, and may thus function as unconventional DCAFs specific for phosphorylated CRL4B.

**Figure 4:**
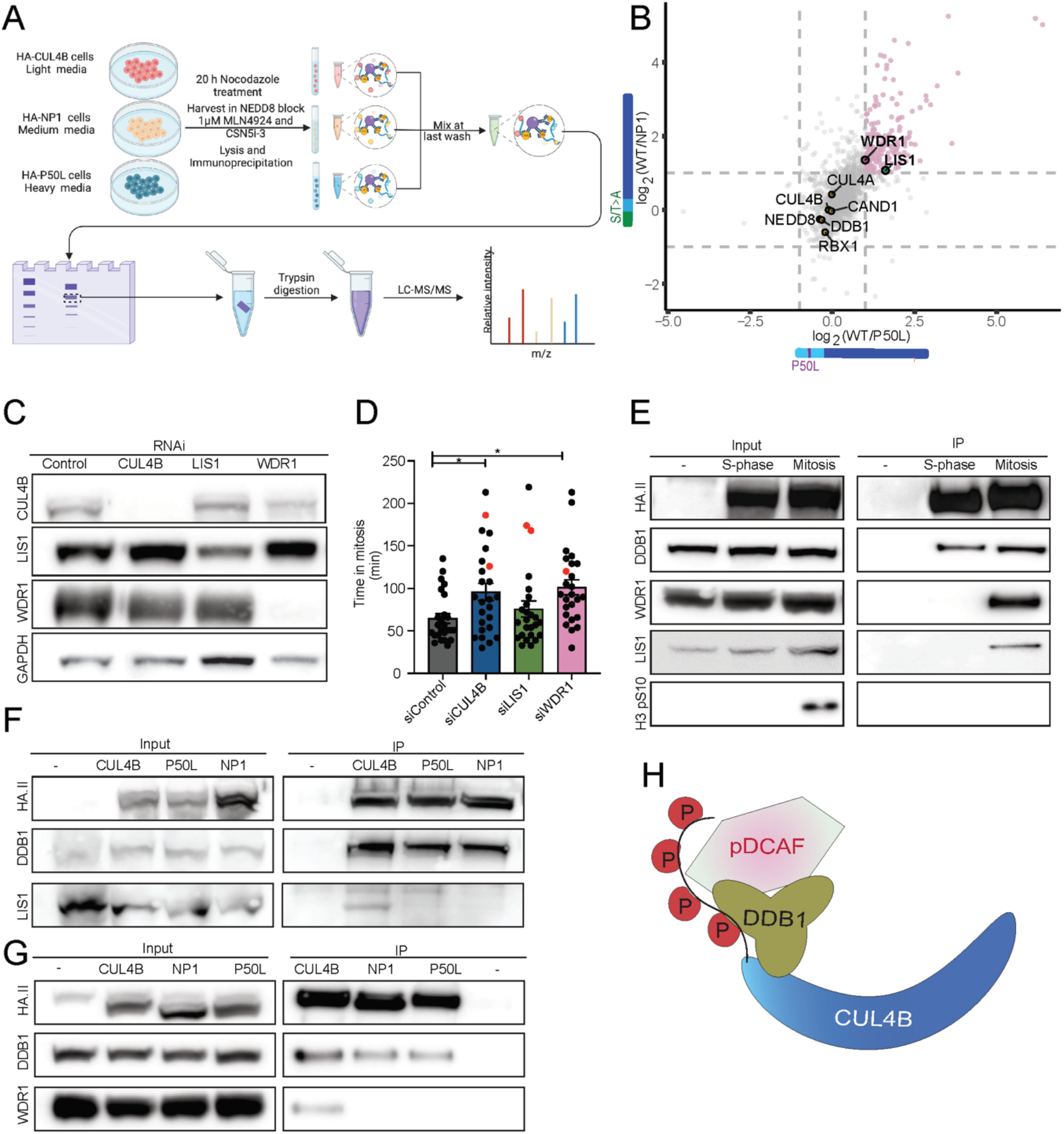
The unconventional DCAFs LIS1 and WDR1 specifically interact with phosphorylated CUL4B during mitosis. (**A**) Outline of the SILAC immunoprecipitation experiment. HeLa Kyoto cells endogenously expressing HA-tagged WT CUL4B or the NP1 or P50L mutants were labeled with heavy, medium or light SILAC medium, and arrested in mitosis with nocodazole for 20 hours. Cell lysis was performed in the presence of the CSN inhibitor CSN5i-3 and NAE inhibitor MLN4924 to prevent DCAF exchange. Samples were mixed after immunoprecipitation with HA.II antibodies and bound proteins separated by SDS-PAGE before trypsin digestion and MS-analysis. (**B**) Scatter plot comparing mitotic immunoprecipitations (IP) of phosphorylated CUL4B with the phospho-mutants NP1 and P50L. The axes represent the log2-transformed fold change of WT CUL4B vs mutant IPs, with dashed lines marking 2x fold change cutoffs. Proteins preferentially binding to phosphorylated CUL4B compared to the phosphorylation-defective mutants, such as WDR1 and LIS1, group in the upper right corner. (**C**) To control RNAi efficiency and specificity, HeLa Kyoto cells were incubated for 72 h with control oligos (control) or RNAi-oligos depleting CUL4B, LIS1 or WDR1. Specific downregulation of the indicated proteins was analyzed by immunoblotting. GAPDH controls for equal loading. (**D**) The mitotic duration of siControl cells or cells RNAi-depleted for CUL4B, LIS1 or WDR1 was quantified by life cell microscopy in the brightfield channel measuring the time from cell rounding to telophase. Mean with SEM are depicted of N= 4 total of 25 cells. Red symbols mark cells with incomplete mitosis during the movie. Statistical analysis was performed with 1-ANOVA and Dunnett’s multiple comparison. (**E)** HA-immunoprecipitation (IP) of extracts (input) prepared from unedited (-) or HA-CUL4B endogenously expressing HeLa Kyoto cell lines arrested in S-phase (thymidine) or mitosis (nocodazole) were probed with antibodies against LIS1, WDR1 and DDB1. The antibody recognizing phospho-H3(S19) controls mitotic arrest and specificity of the immunoprecipitation. (**F**) and (**G**) HeLa Kyoto endogenously expressing HA-CUL4B or the indicated NP1 or P50L mutants were arrested in mitosis with nocodazole. Extracts (input) and HA-immunoprecipitants were analyzed for the presence of HA-CUL4B for control and bound DDB1 and LIS1 (**F**) or DDB1 and WDR1 (**G**), respectively (N=3). (**H**) Schematic model visualizing the assembly of phospho-specific DCAFs (pDCAF)-containing E3 ligase complexes with N-terminally phosphorylated CRL4B during mitosis. See also Extended Data Fig. 4.

To validate these findings, we focused on three main aspects, namely functional overlap with CUL4B during mitosis, as well as mitosis- and phosphorylation-specific binding. To test whether depletion of LIS1 and/or WDR1 can explain the mitotic defects observed in cells lacking CUL4B, we used live-cell imaging to determine mitotic duration. While RNAi-depletion of CUL4B and WDR1 was highly efficient, LIS1 knockdown was moderate (Fig. 4C). Interestingly, WDR1- and to a lesser extend LIS1-depleted cells were delayed in mitosis, thus phenocopying at least in part CUL4B-depletion (Fig. 4D, Extended Data Fig. 4B). The mitosis-specific interaction of CUL4B with LIS1 and WDR1 was verified by immunoprecipitating HA-tagged CUL4B from cells arrested in S-phase by thymidine or in mitosis with nocodazole (Fig. 4E). Phospho-specificity was confirmed by comparing immunoprecipitations of HA-tagged CUL4B and the phosphorylation-defective mutants NP1 and P50L in cells arrested in mitosis by nocodazole. Indeed, these assays demonstrated mitosis-specific binding of CUL4B to LIS1 and WDR1, which was not observed when analyzing the phosphorylation-defective mutants NP1 and P50L (Fig. 4F, G). Together, these results suggest that LIS1 and WDR1 function as phospho-specific DCAFs (pDCAF), assembling with phosphorylated CRL4B (pCRL4B) to regulate progression through mitosis (Fig. 4H).

### LIS1 and WDR1 binding is mediated via two different binding sites one on DDB1 and one on CUL4B

We next used reconstituted complexes (DDB1, RBX1 and either CUL4A or CUL4B) and established a fluorescence polarization (FP)-assay to dissect binding affinities of LIS1 and WDR1 to different CRL4 subunits, Since LIS1 and WDR1 possess the canonical helix-loop-helix motif predicted to mediate DDB1 binding, we performed pull-down experiments using LIS1, WDR1 or for control DCAF8 from Sf9-extracts co-expressing DDB1. Indeed, both LIS1 and WDR1 directly interact with DDB1 under these conditions, although with weaker affinity compared to the conventional DCAF8 (Fig. 5A, Extended Data Fig. 5A). FP assays confirmed that LIS1 and WDR1 bind DDB1 with a Kd of approx. 5 μM (Fig. 5B). Surprisingly, we also detected interaction of LIS1 and WDR1 with CUL4B after co-expression in Sf9 cells in the absence of DDB1. This implies two distinct binding sites of LIS1 and WDR1, one on DDB1 and another one on the N-terminus of CUL4B (Fig. 5 C, D). Consistent with this hypothesis, purified LIS1 and WDR1 preferentially interacted with CRL4B compared to CRL4A, which lacks the N-terminal extension (Fig. 5 E, G; Extended Data Fig. 5B, C). Strikingly, LIS1 and WDR1 exhibit a pronounced binding preference to CRL4B with a Kd around 1 μM, while binding to CRL4A is almost undetectable (Kd around 13 μM) (Fig. 5F, H; Extended Data Fig. 5D).

**Figure 5:**
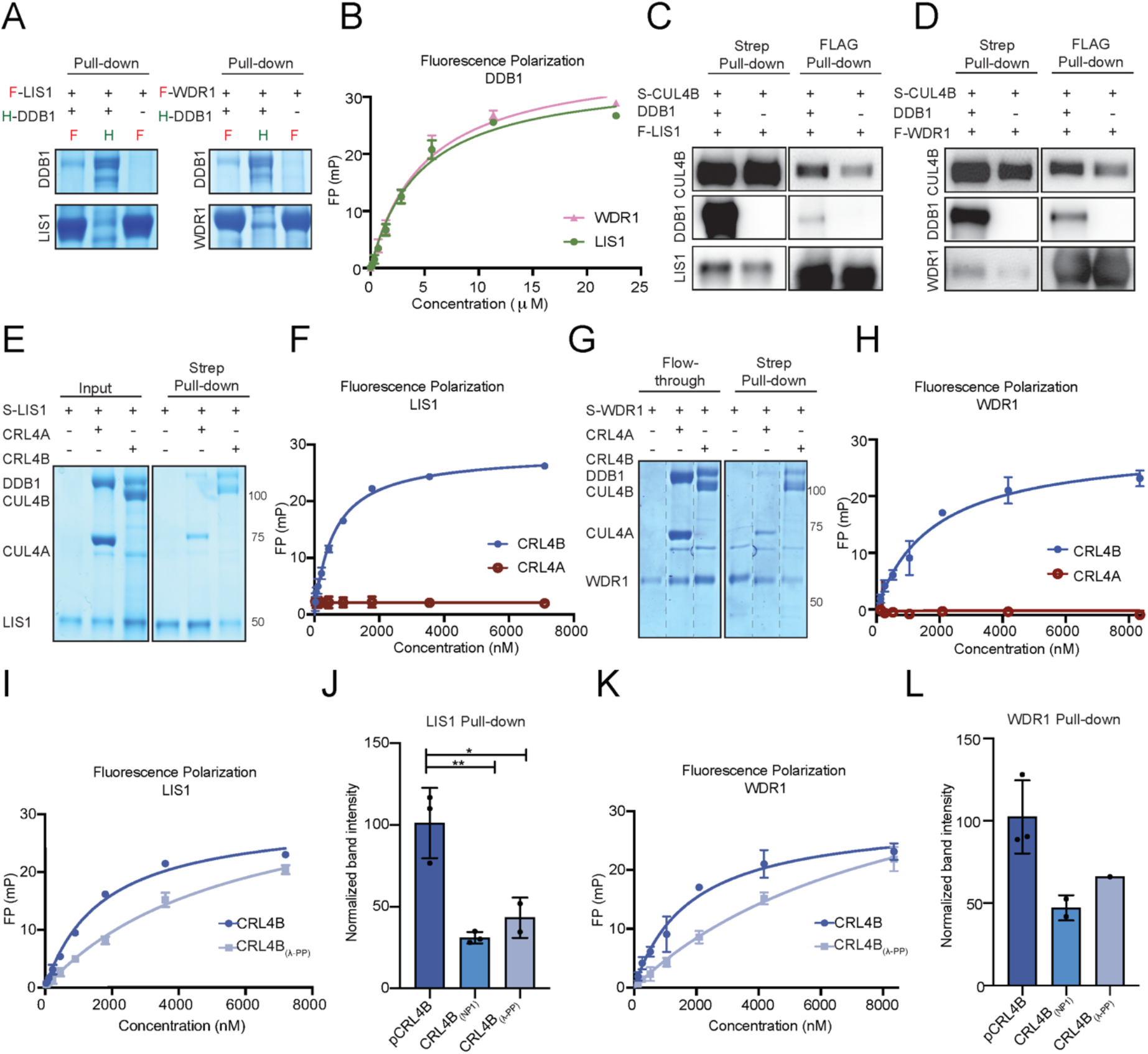
Recruitment of LIS1 and WDR1 into the CRL4B complex is mediated via two different binding sites, one on DDB1 and one on phosphorylated CUL4B. (**A**) Coomassie-stained SDS-PAGE of *in vitro* pull-down assays of FLAG-tagged LIS1 and WDR1, and His-DDB1, co-expressed in baculoviral *Sf9* cells (N=3). For control, binding of Strep-DCAF8 to His-DDB1 was included (N=2). Note that DDB1 directly interacts with LIS1 and WDR1. (**B**) Fluorescence polarization (FP) assay quantifying direct binding of Alexa-labeled LIS1 (N=3) and WDR1 (N=2) to DBB1. (**C**) and (**D**) Western blot of *in vitro* pulldown assays of Strep-tagged CUL4B and Flag-tagged LIS1 (**C**) or WDR1 (**D**) co-expressed in Sf9 cells in the presence (+) or absence (-) of DDB1 (N=3). Note that LIS1 and WDR1 directly bind CUL4B, but this interaction is enhanced by the presence of DDB1. (**E**) Coomassie-stained SDS-PAGE of purified proteins (input) and *in vitro* binding assays (Strep-pull down) of Strep-LIS1 with purified CRL4A or CRL4B (N=3). (**F**) FP assay investigating binding of CRL4B and CRL4A to Alexa-labeled LIS1 (N=3). (**G**) Coomassie-stained SDS-PAGE of purified proteins (input) and *in vitro* binding assays (Strep-pull down) of Strep-WDR1 with purified CRL4A or CRL4B (N=2). (**H**) Fluorescence polarization assay investigating the binding of CRL4B and CRL4A to Alexa-labeled WDR1 (N=3). (**I**) Fluorescence polarization assay comparing the binding of CRL4B and λ-PP-treated CRL4B to Alexa-labeled LIS1. (**J**) Quantification of *in vitro* binding assays (Strep-pull down) of Strep-LIS1 with purified CRL4B, the NP1 phospho-mutant (CRL4B_NP1_), or λ-PP treated (dephosphorylated) CRL4B (CRL4B_λ-pp_). (**K**) Fluorescence polarization assay comparing the binding of CRL4B and λ-PP-treated CRL4B to Alexa-labeled LIS1. (**L**) Quantification of *in vitro* binding assays (Strep-pull down) of Strep-LIS1 with purified CRL4B, or the NP1 phospho-mutant (CRL4B_NP1_), or λ-PP treated CRL4B (CRL4B_λ-pp_). Band intensities were quantified and normalized to LIS1 and WDR1, respectively (N=3 for LIS1; N=2 for WDR1). Statistical analysis: 1-ANOVA followed by Dunnett’s multiple comparison. Note that LIS1 and WDR1 preferentially bind to phosphorylated CRL4B. See also Extended Data Fig. 5.

To investigate whether phosphorylation may enhance the amino-terminal interaction, we first compared CRL4B purified from *Sf9* cells, after additional treatments with either CDK1 or λ-PP. While incubating *Sf9*-expressed CUL4B by CDK1 (CRL4B_CDK1_) did not alter its migration on SDS-PAGE, λ-PP treatment (CRL4B_λ-PP_)led to a faster migrating CUL4B band, similar to the phosphorylation-defective NP1 mutant (CRL4B_NP1_) (Fig. S5E). We thus measured the binding affinity of LIS1 or WDR1 to *Sf9*-purified CRL4B with or without λ-PP treatment. Indeed, dephosphorylation of CRL4B reduced its binding affinity to LIS1 and WDR1 at least 4-fold, reaching the affinity observed by direct DDB1 binding (Fig. 5I, K). To corroborate these data, we also performed *in vitro* pulldown assays of purified LIS1 and WDR1 with CRL4B, CRL4B_NP1_ and CRL4B_λ-PP_. While CRL4B binds LIS1 or WDR1 in an almost stoichiometric ratio, we observed decreased binding of LIS1 and WDR1 to CRL4B_NP1_ or λ-PP -treated CRL4B (CRL4B_λ-PP_) (Fig. 5J, L; Extended Data Fig. 5F, G). Taken together, our results suggest that in addition to DDB1, the phosphorylated N-terminus of CUL4B stabilizes the interaction with the pDCAFs LIS1 and WDR1.

### CRL4B^LIS1^ and CRL4B^WDR1^ complexes regulate cortex tension and spindle positioning during mitosis

Consistent with the reported functions of WDR1 and LIS1 in regulating actin dynamics^33^ and the dynein/dynactin complex^34^, respectively, our phospho-specific, mitotic CUL4B interactome revealed multiple proteins involved in cytoskeletal regulation. Indeed, the GO term analysis of the mass spectrometric dataset found strong enrichment of components involved in cytoskeleton organization and actin filament-based processes (Fig. 6A, B, Table S2 and 3). Interestingly, the Arp2/3 complex was specifically immunoprecipitated by HA-tagged CUL4B in mitosis and its interaction was strongly reduced when analyzing the phosphorylation-defective NP1 and P50L mutants (Fig. 6C). Likewise, HA-CUL4B immunoprecipitates prepared from extracts of mitotic cells (Fig. 6D) contained actin and dynactin, which are known interactors of WDR1 and LIS1, respectively.

**Figure 6:**
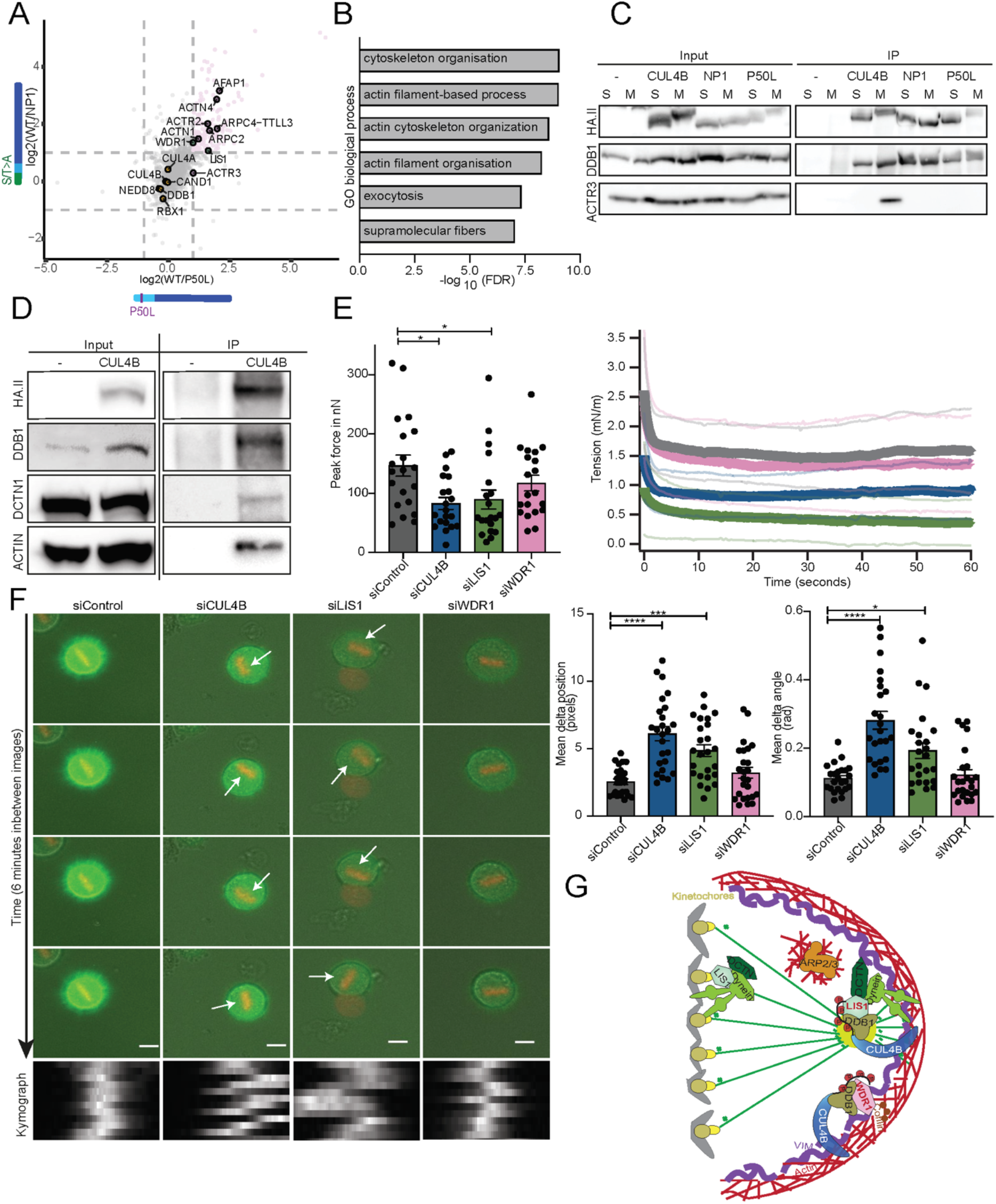
CRL4B^LIS1^ and CRL4B^WDR1^ E3 ligase complexes may regulate cortex tension and spindle positioning during mitosis. (**A**) Mass spectrometric analysis of HA-immunoprecipitants (IP) using extracts prepared from SILAC labeled, nocodazole-arrested (mitotic) HeLa Kyoto cells endogenously expressing HA-tagged WT CUL4B or the NP1 and P50L phospho-mutants. The workflow was described in Figure 4A, and the data displayed as scatter plot. The axes represent the log2-transformed fold change of phosphorylated WT CUL4B vs mutant IPs, with dashed lines marking 2x fold change cutoff. Proteins preferentially binding to phosphorylated CUL4B cluster in the upper right corner. (**B**) GO term biological process analysis of the interactome preferentially binding phosphorylated CUL4B vs the NP1 mutant. (**C**) Extracts (input) prepared from S-phase (S; thymidine) or mitosis (M; nocodazole)-arrested HeLa Kyoto cells (-) and endogenously expressing HA-tagged WT CUL4B, or the NP1 or P50L mutants were immunoprecipitated with HA.II-antibodies (IP). Bound ACTR3 or DDB1 was detected using specific antibodies (N=3). Note that in contrast to DDB1, ACTR3 binds CUL4B in a phosphorylation-dependent manner. (**D**) HA-immunoprecipitants (IP) of extracts (input) prepared from unedited (-) controls or endogenously expressing HA-CUL4B HeLa Kyoto cells arrested in mitosis with nocodazole were probed with antibodies against DDB1, DCTN1 (dynactin) and actin (N=2). (**E**) AFM measurements of the peak confinement force (left) and cortical tension (right) in mitotic cells. HeLa Kyoto cells were incubated for 72 h with control oligos (siControl) or RNAi-oligos depleting CUL4B, LIS1 or WDR1. After 30 min of (+)-S-trityl-l-cysteine (STC) treatment, the confinement force was measured for single mitotic cells by AFM. The initial or peak force is displayed as the mean in nano-Newton (nN) with SEM (N=2, with 10 cells in each experiment) and the tension during confinement in mN m^-1^ for a representative replicate is shown in the right panel. (**F**) Live cell imaging of HeLa cells expressing H2B-mcherry and lifeAct-GFP, treated for 72 h with control oligos (siControl) or RNAi-oligos depleting CUL4B, LIS1 or WDR1. Representative cells are shown in time intervals of 6 min during metaphase with identical contrast settings. The unanchored spindle in CUL4B- and LIS1 deleted cells is marked with a white arrow. Scale bar: 10 μm. The mean delta position (left bar graph) and the mean delta angle (right bar graph) with SEM quantifies the average spindle movement (in pixel per time frame) or spindle rotation (in rad) (N=4, including 25 cells). Only cells with completed mitosis were measured. Statistical analysis was performed by 1-ANOVA with Dunnett’s multiple comparison to RNAi control. The kymographs visualize the spindle movement during metaphase. (**G**) Schematic representation of CRL4B bound to the phospho-specific DCAFs LIS1 and WDR1 in cytoskeletal regulation during mitosis. The left half illustrates CRL4B bound to chromatin in the nucleus during interphase, while the right half depicts cytoskeletal functions of phosphorylated CRL4B in complex with LIS1 and WDR1 during mitosis. The centrosome is shown as yellow circle and chromatin/chromosomes are depicted in grey. Functions of LIS1 and the dynein/dynactin complex at kinetochores and the cell cortex are highlighted. Green lines represent spindle and astral microtubules, red lines cortical actin filaments, possibly regulated by the Arp2/3 complex and WDR1/cofilin activity. Vimentin (Vim) filaments stabilizing the cell cortex are indicated in violet. See also Extended Data Fig. 6.

To functionally dissect possible cytoskeletal functions of pCRL4B^LIS1^ and pCRL4B^WDR1^ during mitosis, we RNAi-depleted CUL4B, LIS1 or WDR1 in HeLa cells expressing LifeAct-GFP and H2B-RFP^35^ and monitored actin dynamics as well as spindle movement and positioning. While we did not detect grossly aberrant actin structures in CUL4B or LIS1 RNAi depleted cells, we observed severe blebbing and cell death in WDR1 depleted cells. We also used atomic force microscopy (AFM) in a cell confinement assay to assess actin-dependent cortical tension and cell stiffness during mitosis (Extended Data Fig. 6A). Interestingly, we detected a strong reduction in the initial confinement force (or peak force) in cells lacking CUL4B (Fig. 6E, Extended Data Fig. 6B). Similarly, WDR1 or LIS1 depletion reduced this peak force, although to a lower extent (Fig. 6E). In addition, the tension of the actin cortex was strongly reduced in LIS1 and CUL4B depleted cells, while WDR1 depletion only led to a small decrease (Fig. 6E). Together, these data indicate that CRL4B activity is required to stabilize the cell cortex during mitosis, most likely by regulating actin-dependent structures via LIS1 and WDR1.

Since LIS1 is known to regulate dynein/dynactin complexes and spindle positioning^36^, we tested whether CUL4B is required for proper spindle organization and function. Indeed, immunofluorescence experiments revealed that ΔCUL4B cells show an increased number of off-centered mitotic spindles (Extended Data Fig. 6C). To corroborate these findings, we used live-cell microscopy to follow spindle dynamics in single cells. Kymographs showed enhanced spindle mobility in CUL4B depleted cells and spindle movement increased 5-fold compared to controls. Assessment of the spindle angle indicated that the mitotic spindle was not stably anchored to the cortex with an increased mobility in all three dimensions (Fig. 6F). As expected, similar spindle defects were detected in LIS1-depleted cells, while lack of WDR1 did not significantly alter spindle dynamics (Fig. 6F). Thus, phenotypic characterization indicates that CRL4B^WDR1^ regulates mitotic duration possibly by actin rearrangements, while CRL4B^LIS1^ is required to stably anchor the mitotic spindle to the cell cortex and increase cortex tension (Fig. 6G).

## Discussion

Our study establishes a novel CRL4B-specific regulatory mechanism comprising cell-cycle dependent phosphorylation of its unique unstructured N-terminal domain. We demonstrate that phosphorylated CUL4B engages with the previously unrecognized DCAFs, LIS1 and WDR1, which together regulate cortex tension and spindle positioning during mitosis. Since the P50L patient mutation is specifically defective for its mitotic phosphorylation and in turn fails to bind LIS1 and WDR1, our findings provide important insights to understand the molecular mechanisms causing XLID and possibly other human neurological diseases.

We found that the unique CUL4B-extension contains many S/T-residues, which are phosphorylated during mitosis. Indeed, several mitotic kinases including CDK1, PLK1 or Aurora B phosphorylate CUL4B *in vitro*. CDK1-cyclin B emerged as the major CUL4B kinase preferentially phosphorylating T49, the major site conforming to the minimal S/T-P-X motif^37, 38^. Strikingly, the P50L XLID patient mutation interferes with phosphorylation of the adjacent T49 residue, explaining its mitotic defect. CDK1 sites often promote PLK1 binding, which then further increases phosphorylation^39, 40^. It will be interesting to explore whether non-mitotic kinases may similarly shape appropriate cellular responses by regulating CUL4B function through phosphorylation of its unique amino-terminal domain in response to different environmental conditions.

We found that CUL4B phosphorylation is required for efficient progression through mitosis by two mechanisms. First, mitotic phosphorylation triggers its dissociation from mitotic chromatin, and second, phosphorylation allows binding of CUL4B to previously unrecognized DCAFs, including WDR1 and LIS1. Both WDR1 and LIS1 possess WD40 domains and bind DDB1 most likely through a predicted helix-turn-helix motif^9, 32^. Importantly, however, their interaction is stabilized by a separate binding motif generated by the phosphorylated amino-terminal extension of CUL4B. Together, the two distinct binding sites promote the assembly of stoichiometric CRL4B complexes with low micromolar affinity like conventional DCAFs binding to CRL4s. Biophysical and structural work will be required to understand the underlying properties and deduce critical residues stabilizing the DDB1-LIS1 and DDB1-WDR1 heterodimers bound to phosphorylated CUL4B. It is possible that additional CRL4B-specific DCAFs exist, including unconventional DCAFs which require other modifications of the amino-terminus. An interesting candidate could be AMBRA1, which was recently shown to regulate cyclin D degradation via CUL4B^41–43^. Importantly, functional importance of the unique N-terminal region of CUL4B was also demonstrated for the Dioxin receptor, which binds independently of DDB1^44^. Moreover, WDR1 was previously reported to function as a DCAF in *Trypanosoma brucei*, which encodes CUL4 with an amino-terminal extension like CUL4B^45^. It will thus be interesting to compare evolutionary conserved features, as the unique amino-terminal extension increases the repertoire of substrate receptors specifically functioning within CRL4B complexes. Our observations may also be relevant for targeted protein degradation, since several degraders rely on substrate recruitment by the CRL4^CRBN^ E3 ligase^46^. Exploiting phospho-dependent CRL4B substrate receptors may restrict neo-substrate targeting to proliferating cells in mitosis.

Several studies have addressed molecular functions of LIS1 and WDR1 in mitosis. LIS1 regulates spindle mobility and orientation by regulating the dynein/dynactin complex and by increasing the density of astral microtubules, which may in turn affect cortical tension^36, 47, 48^. Cells lacking CUL4B similarly exhibit spindle positioning and cortical tension defects, and it is thus tempting to speculate that pCRL4B^LIS1^ regulates dynein/dynactin activity. Less is known about the molecular functions of WDR1, but available data suggest that it likely controls the actomyosin network and functions in mitotic cell rounding^49, 50^.

While LIS1 and WDR1 coordinately regulate cortical tension and cytoskeletal dynamics, it remains to be examined if and how these cortical defects alter mitotic timing. In mitosis, the cortex becomes thinner with increased tension due to RhoA activation, which allows cells to change to the rounded-up morphology^51^. Both spindle orientation and chromosome separation rely on the actomyosin cortex providing a rigid scaffold to counteract the traction forces exerted on astral microtubules by motors pulling towards the spindle poles^52^. Given the overlapping defects of cells lacking CUL4B, LIS1 and WDR1 in cortical tension and cytoskeletal dynamics, the formation of functional CRL4B^LIS1^ and CRL4B^WDR1^ E3 ligase complexes may contribute to these functions during mitosis. A critical next step to test this exciting hypothesis will be the identification of relevant substrates of mitotic CRL4B complexes. Candidates include proteins directly binding to LIS1 and WDR1, and we indeed found dynactin and many actin-regulators as mitotic-specific interactors of CUL4B. Further studies are needed to demonstrate if these candidates are indeed critical substrates for pCUL4B-dependent ubiquitination.

Our results provide important molecular insight into the function of CRL4B in XLID and related intellectual disability disorders. In contrast to most *CUL4B* patient mutations, the amino-terminal P50L XLID patient mutation neither impacts assembly nor catalytic activity of CRL4B-based E3 ligase complexes *in vitro*. Instead, we demonstrate that this mutation affects CUL4B’s mitotic function by reducing mitotic T49 phosphorylation, thereby preventing its interaction with phospho-specific DCAFs such as LIS1 and WDR1. This regulatory mechanism could explain why CRL4A fails to compensate CRL4B-specific functions causing XLID. Consistent with CRL4B regulating cytoskeletal dynamics, comprehensive analysis reveals that intellectual disability genes converge onto a few biological modules, including cytoskeleton alterations that impact neuronal migration, neurogenesis, and synaptic plasticity^53, 54^. Interestingly, LIS1 mutations are strongly linked to classical lissencephaly (LIS), a neuronal disorder characterized by brain malformation and intellectual disability caused by defects in neuronal migration^55^. Likewise, patients with mutations in WDR1 exhibit a mild form of intellectual disability, but also suffer from aberrant lymphoid immunity^56^. Thus, further research on CUL4B specific functions might elucidate new insights in brain development and dysfunction.

## Acknowledgments

We would like to thank B. Baum for kindly providing the HeLa Kyoto H2B-mcherry and lifeAct-GFP cells. We thank ScopeM and especially S.-S. Lee for microscopy training and support, F. Lampert for help with the CRL4B^CRBN^ ubiquitination assays, and J. Chen and the Proteomics Core Facility at IMB Mainz for assisting the mass spectrometry analysis. We thank K. Reichermeier and D. Kirkpatrick, as well as R. Aebersold for initial mass spectrometry experiments and advice. We are grateful to U. Kutay, P. Meraldi, N. Thomä and members of the Peter lab for helpful discussions, and Alicia Smith for critical editing. Work in the Peter laboratory is supported by the Swiss National Science Foundation (310030_179283/1), the Swiss Cancer league and ETH Zürich. Work in the Beli lab is funded by the Deutsche Forschungsgemeinschaft (DFG, German Research Foundation) – Project-ID BE 5342/2-1 – FOR 2800. Open access funding is provided by ETH Zurich.

## Author Contributions

Conceptualization and Methodology: AS, SG, WIM, TS and MP; Investigation and Validation: AS, SG, WIM, JH and IM. Software and Formal Analysis: JH and IM. Writing-Original draft: AS and MP. Writing-Review and Editing: AS, MP, WIM, with input from all authors. Supervision: MP, PB, DJM. Funding Acquisition: PB, MP.

## Methods

### Antibodies and reagents

For western blot and immunofluorescence analysis following primary antibodies were used: Anti-Actin (mouse, Merck Millipore); Anti-ARP3 (mouse, Santa Cruz); Anti-CK1α (mouse, Santa Cruz); Anti-CUL4A (rabbit); Anti-CUL4-CT (rabbit) (Olma et al., 2009); Anti-CUL4B (rabbit, Sigma); Anti-DCTN1 (rabbit, Atlas antibodies); Anti-DDB1 (mouse, BD Biosciences); Anti-GAPDH (mouse, Sigma); Anti-HA.II (mouse or rabbit, both Covance); Anti-Histone 4 (rabbit, Abcam); Anti-Phospho-Histone H2A.X S139 (mouse, Merck Millipore); Anti-Phospho-Histone H3 S10 (rabbit, Upstate); Anti-LIS1 (mouse, Santa Cruz Biotechnology); Anti-Pericentrin (rabbit, Covance); Anti-Tubulin (mouse, Sigma); Anti-WDR1 (rabbit, Atlas antibodies). For western blot Anti-mouse or rabbit IgG-HRP (both Biorad) and for immunofluorescence anti-mouse or rabbit IgG-Alexa 488, 568 and 647 (Invitrogen) secondary antibodies were used. Rhodamine phalloidin dye was used for acting staining.

### Cell culture, RNAi and transient transfection

Cells were cultured at 37°C with 5% CO2 in DMEM (Gibco) supplemented with 10% FCS and 1% penicillin-streptomycin-glutamine (Gibco).

RNAi experiments were performed according to manufacturer’s instructions with 25 nM double stranded siRNA and Lipofectamine RNAiMax (Invitrogen). Depending on the experiments the cells were harvest after 48 to 96 h after RNAi. siRNAs (Microsynth) used: siCUL4A (5’-GAATCCTACTGTTGATCGATT), siCUL4B (5’-AGCGCCTGTTAGTCGGAAA), siDDB1 (5’-ACTAGATCGCGATAAATT), siLIS1 (5’-CGGACAAGTAGAATAAATGTT); siWDR1 (5’-GGTGGGATTTAGGCAATTATT) and AllStars Negative Control siRNA (Qiagen).

Transient transfection was accomplished using lipofectamine 3000 (Invitrogen) and FLAG-CUL4B and mutants in pcDNA5/FRT/TO vector (Invitrogen).

### Generation of stable cell lines

The CUL4B knock-out cell lines were achieved with SpCas9-mediated genome editing based on the protocol of Ran et al 2013 ^57, 58^. The gRNAs targeting the intron between exons 2 and 3 (target DNA: 5’TCCCGCGGGACCGTTAAGGA) and between exons 4 and 5 (target DNA: 5’TAACGGCTACCTATATGGTA) were cloned into a pX458 plasmid, kindly provided by F. Zhang (Addgene plasmid #48138) ^59^ and were transient transfected using lipofectamine 3000 transfection reagent (Invitrogen). GFP positive cell sorting in a BD FACSAria IIIu cell sorter (BD Biosciences) was performed four days after transfection. Individual clones were picked and tested for CUL4B deletion by western blot. Knock-in of HA-tagged CUL4B or mutants in the CUL4B gene locus was achieved by targeting two regions within exon 3 (target DNA 1: GGTGCTGGTATTACCATCAG; target DNA 2: ACTAACTTCTTAGCAGAGCC). The pX458 plasmids containing the gRNAs were co-transfected with pUC18 plasmid, the Puro-P2A-HA containing template vector for homologous recombination. Four days after transfection cells were selected with 1 μg/mL puromycin. Individual clones were picked and tested by western blot.

Generation of Flp-In T-Rex HeLa cell lines with inducible expression of CUL4B or mutants was accomplished with co-transfection of pcDNA5-pDEST-FRT-CUL4B/ mutants and the pOG44 vectors adapted from Kean et al. 2012 ^60^. Cells were selected 24 h after transfection with 200 μg/mL Hygromycin B and single colonies were picked and tested for Doxycycline-inducible expression of CUL4B and mutants.

### MTT assay

24 h after RNAi 2,5 x10^3^ cells per well were seeded in 4 x 96 well plates with 3 technical replicates per condition. The MTT assay kit (Promega) was used according to manufacturer’s instructions. The MTT dye master mix was incubated for 2 h at 37°C and 5% CO2 on the cells and the reaction was stopped with the addition of 100 μl stop solution. The plate was measured at a wavelength of 570 nm and the growth rate was calculated normalizing to day 0.

### SDS-PAGE and Western blot

Samples were either run on 4-12% NuPAGE or Bolt gels with the NuPAGE Bis-Tris system (Invitrogen) in MOPS SDS buffer or on 8% gels with the BIO-RAD system with SDS-running buffer. Western blotting was performed with the wet transfer systems from BIO-RAD on 0.45 μm Nitrocellulose membranes (Amersham™), which were blocked in 5% milk in PBS. Primary antibody dilutions were 1:500 in 5% milk-PBS, while secondary antibodies were diluted 1:2000. Blots were visualized with SuperSignal™ West Pico PLUS or Femto Chemiluminescent Substrate solution (Thermo Fischer) or with Clarity™ Western ECL solution (BIO-RAD) and scanned on Fusion FX7 imaging system (Witec AG).

### Immunofluorescence

Cells were seeded on coverslips one or two days before fixation with 4% PFA for 10 min. For cytoplasmic pre-extraction cells were incubated with extraction buffer (20 mM PIPES pH 6.8, 10 mM EGTA, 1 mM MgCl2 and 0.25% Triton X-100) for 4 min before fixation. Cells were permeabilizated with 0.25% Triton X-100 in PBS and washed with PBS with 0.05% Triton X-100. Blocking buffer (PBS, 0.05% Triton X-100, 5% FCS) was used for 30 min before the primary antibody dilution 1:500, which was incubated for 1 h at RT. The samples were washed 3 times and incubated for 1 h with 1:4000 secondary anti-mouse IgG and anti-rabbit IgG antibodies, conjugated to Alexa 488, Alexa 568 or Alexa 647. After three washes the coverslips were mounted on, slides with Immu-mount (Thermo Scientific). The second wash was supplemented with 0.1 μg/mL DAPI (Sigma) and for cytoskeletal staining phalloidin dye. For DNA damage assessment cells were imaged in a single plan with a 20x objective, while for chromatin exclusion and spindle positioning Z-stacks of cells were acquired in 0.33 μm steps from 5 μm to −5 μm in an Eclipse Ti epifluorescence microscope (Nikon) using a 60x oil objective (NA 1.4 Plan Apochromat). Example images for spindle positioning were taken with the Olympus Fluoview 3000. Images analysis was performed with ImageJ converting Z-stacks to maximum projections using the same contrast settings of per channel for all the conditions. Nuclear intensity was measured with a mask based on DAPI signal, while spindle positioning was measured based on the pericentrin signal localization in regard to the center of the cells.

### Live-cell imaging

For mitotic duration assessment around 10 000 cells were re-seeded in 8-well μ-Slide chambers (Ibidi) after 48 h of siRNA treatment or 24 h of Doxycycline induction in 6 well plates. Cells were imaged every 3 min in the brightfield channel (BF) on the Eclipse Ti epifluorescence microscope (Nikon Instruments), at 37°C and 5% CO2 and stable humidity, using a 10x objective (NA 0.45 CFI Plan Apochromat).

For mitotic duration, approximately 10 cells per movie were analyzed counting the frames from rounding of the cell to cytokinesis of a single cell.

The spindle movement was assessed with the HeLa Kyoto H2B-mcherry and lifeAct-GFP cells, which were a kind gift from B. Baum^35^. Imaging was performed as described above every 3 min in BF, RFP and GFP channels with 5% LED intensity. To track the position of the mitotic plate an in-house IGOR (WaveMatrics) code was used. Only cells that both entered and completed mitosis during each time-lapse recording were analyzed. In each image the spindle was found using a threshold of which the position and angle were determined by fitting an ellipse.

### Cell cycle arrests

Cell cycle arrests were performed with different drugs dependent on cell cycle phase. Cells were arrested in S-phase with 2 mM thymidine (Sigma), in G2 with 10 μM RO3306 (Calbiochem) and in mitosis with 100 ng/μL nocodazole (Sigma) or taxol (Paclitaxel, Sigma) for 24 h. Mitotic arrest for AFM experiments were performed with 2 μM (+)-*S*-trityl-L-cysteine (STC, Sigma) for 1 h.

### G2 block and release

Cells were seeded in 6 well plates and arrest in G2 for 20 h with 10 μM RO3306 (Calbiochem) addition. Release was performed with 2x PBS washes and 1 media wash. Synchronized cells were harvested at different time points after the release in 1x Leammli buffer.

### λ-Phosphatase treatment

4 x 10^6^ cells were lysed in 1x PMP buffer supplemented with 1 mM MnCl_2_, 0.5 % NP40, 0.5 mM PMSF and protease inhibitor cocktail. Lysates were split in two tubes and either 200 U λ-Phosphatase or lysis buffer were added, followed by an incubation for 1 h at 30°C.

### Protein expression and purification

cDNAs encoding human CUL4B (Q13620), DDB1 (Q16531), WDR1 (O75083), and LIS1 (P43034) were cloned into pAC8 vector, which is derived from the pBacPAK8 system (ClonTech). CUL4A (FL) (Q13619) and DDB1 (FL) (Q16531) were assembled into the same vector (pBIG1a) using Gibson-based technique (biGBac)^61, 62^. FLAG-tagged CUL4, DDB1 or RBX1 constructs were also cloned into pFL multiBac vectors. Bacmids were produced in DH10Bac/Multibac E. coli bacterial strains and extracted as previously described^63^. Recombinant baculoviruses were prepared in *Spodoptera frugiperda* (*Sf9*) cells using Cellfectin (Invitrogen) following the Bac-to-Bac protocol (Life Technologies). Recombinant protein complexes were expressed in *Sf9* or *Trichoplusia ni* High Five cells by co-infection of single baculoviruses. For the CRL4B WT or NP1 complexes (CUL4B, DDB1, and Rbx1), the CUL4B (WT or NP1 mutant) was expressed with N-terminal Strep (II) tag and DDB1 and RBX1 with N-terminal His-tag. For the CRL4A complex (CUL4A, DDB1, and Rbx1), all components were expressed with an N-terminal His-tag. WDR1 and LIS1 were expressed separately, each with an N-terminal Strep (II) tag, or Strep II-Avi tag. Cells were harvested 48 h after infection and lysed by sonication in a buffer containing Tris-HCl pH 8.0, 200 mM NaCl, and 0.5 mM TCEP, including 0.1% Triton X-100, 1x protease inhibitor cocktail (Roche Applied Science) and 1 mM phenylmethanesulfonyl fluoride (PMSF). Lysates were cleared by ultracentrifugation for 45 min at 40,000 *g*. The supernatant was loaded on Strep-Tactin (IBA life sciences) or Ni-NTA (Sigma) affinity chromatography resins in a buffer containing Tris-HCl pH 7.5, 200 mM NaCl and 0.5 mM TCEP. The Strep (II) and Ni-NTA elution fractions were further purified via ion exchange chromatography (Poros HQ 50 μm, Life Technologies) and subjected to size-exclusion chromatography in a buffer containing 50 mM HEPES pH 7.4, 200 mM NaCl, 0.5 mM TCEP, and 10% glycerol. Pure fractions, as judged by SDS-PAGE, were collected, and concentrated using 10,000 MWT cut-off centrifugal devices (Amicon Ultra) and stored at −80 °C. For FLAG-tagged CUL4A/B, DDB1, RBX1 complex, infected cells were resuspended lysed in buffer containing 50 mM Tris-HCl pH 8.0, 350 mM NaCl, 0.4% Triton X-100, 1 mM EDTA, 1 mM DTT and protease inhibitor cocktail (Roche), using a Dounce homogenizer. Lysates was cleared by centrifugation at 12 000 g for 30 min at 4°C and loaded into anti-FLAG M2 affinity gel (Sigma). Washes were performed in buffer containing 50 mM Tris-HCl pH 8.0 and 150 mM NaCl and CRL4 complexes were subsequently eluted with 3xFLAG peptide diluted in 50 mM Tris-HCl pH 8.0, 150 mM NaCl and 10% glycerol, frozen in liquid nitrogen and stored at −80°C until analysis.

### In vitro kinase assay

0.5 μg of recombinant FLAG-CUL4A or FLAG-CUL4B (in complex with DDB1 and RBX1) or 0.15 μg native swine myelin basic protein (MBP; SignalChem) were added to 1x protein kinase buffer (NEB) supplemented with 200 μM “cold” ATP (Sigma) and 500 μCi/μmol ATPγ^32^P (Hartmann Analytic). The reaction was initiated by addition of 10 U (pmol/min) of recombinant CK1 (NEB), CDK1-cyclin B1 (NEB), PLK1 (SignalChem), Aurora A (SignalChem) or Aurora B (SignalChem) kinases, incubated for 1h at 30°C. Samples were boiled in 1x Laemmli buffer and subjected to SDS-PAGE. Gels were stained with InstantBlue coomassie stain (Expedeon Protein Solutions), vacuum-dried for 30 min and exposed overnight to a storage phosphor screen (GE Healthcare). Screens were scanned in a Typhoon FLA 9000 system (GE Healthcare).

### In vitro ubiquitination assay

CRL4B^CRBN^ complexes were assembled by mixing CRL4B WT, NP1 or P50L mutants at 117.5 nM with 12.5 nM DDB1-CRBN in the presence of 10 μM Lenalidomide. *In vitro* ubiquitination of the substrate CK1α was performed by mixing the assembled CRL4B^CRBN^ complexes with a reaction mixture containing UBE1 at 1 μM, UBCH5a at 0.5 μM, Ubiquitin at 15 μM, and 0.25 μM CK1α, as indicated. Reactions were carried out in 50 mM Tris-HCl pH 7.6, 3 mM ATP, 0.5 mM DTT, 10 mM MgCl_2_, and incubated for 0–10 min at 37 °C.

### In vitro pull-down assays

For pull-down assays in *Sf9 cells*, 150 μl of baculoviruses of His-DDB1 and FLAG-WDR1 or LIS1, or Strep-DCAF8 and Strep-CUL4B were co-infected in 10 ml of *Sf9* cells. Infected cells were incubated at 27 °C for 48 h and lysed by sonication in a buffer containing Tris-HCl pH 8, 200 mM NaCl and 0.5 mM TCEP, including 0.1% Triton X-100, 1x protease inhibitor cocktail (Roche Applied Science), 1 mM PMSF, and 10% glycerol. Lysates were cleared by centrifugation at 14,000 *g* for 30 minutes, and 1 ml of soluble protein fractions were incubated for 1 h at 4 °C with 20-30 μl Strep-Tactin Macroprep beads (IBA lifesciences), FLAG or Ni-NTA beads (Sigma). Beads were washed three times with lysis buffer, and bound proteins were eluted in 20-30 μl of SDS loading dye and heated at 95 °C for 2 min.

For pull-down assays using purified protein complexes, Strep-tag was cut out of the CRL4B WT and mutant, and CRL4A complexes by the His-tagged TEV-protease. TEV was then removed by loading the cleaved sample on Ni-NTA beads (Sigma). CDK1 phosphorylation was performed at 30°C for 2.5 h in 1x PK buffer, ATP (2 mM), CRL4B (14 μM) and 19,8 μl CDK1-cyclin B (20000 U/ml). Dephosphorylation of CRL4B took place in a 100 μl reaction mixture containing 86 μl of CRL4B protein (at 0.8 mg/ml) with 2 μl λ-phosphatase (λ-PP) in the presence of 0.1 mM MnCl_2_ and 1x PMP reaction buffer (NEB). Untagged-CRL4 complexes (4A, 4B WT, 4B NP1, and λ-PP-treated 4B) were incubated with Strep-tagged LIS1 or WDR1 at 1:1 molar ration of 1.5 μM in 35 μl. Reaction mixture was loaded on 5 μl Strep-resin (IBA bioscience) and shaking at room temperature. Beads were washed 4 times with 500 μl of buffer containing HEPES pH 7.2 50 mM, NaCl 200 mM, and TCEP 0.5 mM, followed by the addition of 15 μl SDS loading dye on the washed beads before loading 10 μl on NuPAGE (Thermo Fischer). Quantification of band intensities was carried out in ImageJ. CUL4 band intensities were normalized to the band intensity of LIS1 or WDR1.

### Biotinylation

Purified StrepII-Avi-tagged LIS1 or pII-Avi-tagged WDR1 were biotinylated *in vitro* at a concentration of 50 and 38 μM, respectively. Reaction was incubated with 2.5 μM BirA enzyme and 0.2 mM D-Biotin in 50 mM HEPES pH 7.4, 200 mM NaCl, 10 mM MgCl2, 0.5 mM TCEP and 20 mM ATP. The reaction was incubated for 3 h at room temperature and at 4 °C overnight. Biotinylated proteins were purified by size exclusion chromatography and stored at −80 °C.

### Fluorescence Polarization

Alexa488-coupled streptavidin (Thermo Fischer) were incubated at 20 μM in 1:1 molar ratio with biotinylated LIS1 (20 μM) or WDR1 (18 μM) for 30-60 minutes at room temperature and buffer exchange (Zeba Spin) was performed in the reaction buffer (HEPES pH 7.4, NaCl 200 mM, D-Trehalose 25 mM, Triton X-100 0.01%, TCEP 0.5 mM, and BSA 0.1 mg/ml. 26 nM of Alexa-labeled LIS1, or 15 nM WDR1 were titrated with increasing concentrations of DDB1, CRL4B or CRL4A as indicated. Reactions took place at room temperature in 384-well plates in a BMG Clariostar plate reader (BMG LabTech). Binding affinities were calculated by measuring the change in fluorescence polarization. Data were plotted and analyzed using GraphPad Prism assuming a single site binding saturation.

### Cell fractionation

The NE-PER Nuclear and Cytoplasmic Extraction Reagents (Thermo Fischer) were used according to the manufacturer’s instructions to isolate the nuclei. 48 h after transfection around 4 million cells were harvested by trypsinization and nuclei were isolated and cytoplasmic samples were taken. The isolated nuclei were lysed in a buffer containing 10 mM Hepes-KOH pH 7.7, 100 mM KCl, 50 μM sucrose, 0.25% Triton X-100 and protease inhibitor cocktail with a 27G syringe. Nuclear lysates were then loaded on top of a 1 M sucrose cushion and centrifuged at 8000 g for 30 min. To separate the pellet (DNA bound) and the soluble fraction the lysates were loaded on a 1 M sucrose cushion and centrifuged for 30 min at 8000 g. Samples were boiled in 1x Laemmli buffer for 5 min at 95 °C followed by SDS-PAGE and western blot.

### Immunoprecipitation

Around 4 x 10^7^ cells expressing HA-CUL4B or mutants were harvested in a NEDD8 block ^19^. Thus 1 μM MLN4924 and CSN5i-3 was added to the PBS wash and to the lysis buffer (IP buffer plus DTT). Lysis was performed for 40 min using a 27G needle syringe and with the addition of 125 U pierce universal nuclease (Thermo Fischer). The IP buffer contained 20 mM HEPES pH 7.9, 100 mM KCl, 2 mM MgCl_2_, 0.5% NP40, 300 mM sucrose, 10 mM NaF, 10 mM β-glycerophosphate, 0.2 mM NaVO4 and 1 mM PMSF. After centrifugation for 30 min at 16000 g, protein concentration was measured with Quick Start Bradford reagent (Bio-Rad) and normalized to lowest sample. Samples were incubated with anti-HA-agarose beads (Sigma; clone HA-7) for 1 h at 4°C. The beads were washed 5x with IP buffer. Proteins were eluted with 1x Laemmli buffer and analyzed on SDS-PAGE followed by western blot.

### Profiling of CUL4B phosphorylation sites by quantitative mass spectrometry

HA-tagged CUL4B was immunoprecipitated form around 1×10^8^ mitotic cells as described above. After a last wash with MilliQ water the dried beads were frozen in liquid nitrogen and stored at −80°C. The proteins were eluted from beads by 3×30 min incubation in 6 M urea / 2 M thiourea at room temperature. Supernatants were reduced with 1 mM DTT and alkylated with 5.5 mM chloroacetamide. Proteins were pre-digested with 1:100 w/w LysC (Wako) for 4 h, diluted in 4 volumes of water and digested with 1:100 w/w trypsin (Serva) overnight. Samples were incubated in TFA (0.5% V/V) for 1 h at 4°C and centrifuged for 10 min at 4000 × g. Peptide supernatants were purified using C18 Sep-Pak columns (Waters), eluted in 50% acetonitrile and acidified with TFA to 6% V/V. Phosphopeptides were enriched using titanium dioxide resin as described^64^ and desalted using reversed-phase C18 StageTips^65^.

Samples were analyzed on a quadrupole Orbitrap mass spectrometer (Exploris 480, Thermo Scientific) equipped with a UHPLC system (EASY-nLC 1200, Thermo Scientific). They were loaded onto a C18 reversed-phase column (55 cm length, 75 mm inner diameter) and eluted with a gradient from 2.4 to 32% ACN containing 0.1% formic acid in 90 min. The mass spectrometer was operated in data-dependent mode, automatically switching between MS and MS2 acquisition. Survey full scan MS spectra (m/z 300–1,650, resolution: 60,000, target value: 3e6, maximum injection time: 60 ms) were acquired in the Orbitrap. The 15 most intense precursor ions were sequentially isolated, fragmented by higher energy C-trap dissociation (HCD) and scanned in the Orbitrap mass analyzer (normalized collision energy: 30%, resolution: 30,000, target value: 1e5, maximum injection time: 60 ms, isolation window: 1.4 m/z). Precursor ions with unassigned charge states, as well as with charge states of +1 or higher than +6, were excluded from fragmentation. Precursor ions already selected for fragmentation were dynamically excluded for 25 s. Raw data files were analyzed using MaxQuant (version 1.5.2.8)^66^. Site localization probabilities were determined by MaxQuant using the posttranslational modification scoring algorithm. Parent ion and MS2 spectra were searched against a reference proteome database containing human protein sequences obtained from UniProtKB (version 2020_02) using Andromeda search engine ^67^. Spectra were searched with a mass tolerance of 6 p.p.m. in MS mode, 20 p.p.m. in HCD MS2 mode, strict trypsin specificity, and allowing up to two miscleavages. Cysteine carbamidomethylation was searched as a fixed modification, whereas protein N-terminal acetylation, methionine oxidation, phosphorylation (STY) and N-ethylmaleimide modification of cysteines (mass difference to cysteine carbamidomethylation) were searched as variable modifications. The dataset was filtered based on posterior error probability (PEP) to arrive at a false discovery rate of below 1% estimated using a target-decoy approach^68^.

### CUL4B interaction profiling by quantitative mass spectrometry

HA-tagged CUL4B or the phosphorylation deficient mutants NP1 and P50L were immunoprecipitated as described above from around 7,5 x 10^7^ SILAC labeled and nocodazole arrested cells (mitotic). During the last wash, the beads from all three conditions were combined. Bound proteins were eluted in 2x NuPAGE LDS Sample Buffer (Life Technologies) supplemented with 1 mM DTT, heated at 70 °C for 10 min, alkylated by addition of 5.5 mM chloroacetamide for 30 min, and separated by SDS–PAGE on a 4–12% gradient Bis–Tris gel. Proteins were stained using the Colloidal Blue Staining Kit (Life Technologies) and digested in-gel using trypsin (Serva). Peptides were extracted from the gel using a series of increasing acetonitrile percentages and desalted using reversed-phase C18 StageTips. Mass spectrometry acquisition and analysis were performed as described above with following modifications. Peptide fractions were analyzed on a quadrupole Orbitrap mass spectrometer (Q Exactive Plus, Thermo Scientific) equipped with a UHPLC system (EASY-nLC 1000, Thermo Scientific). Peptide samples were loaded onto C18 reversed-phase columns (23 cm length, 75 μm inner diameter, 1.9 μm bead size) and eluted with a linear gradient from 1.6 to 52 % acetonitrile containing 0.1% formic acid in 175 min. The mass spectrometer was operated in a data-dependent mode, automatically switching between MS and MS2 acquisition. Survey full scan MS spectra (m/z 300–1,650, resolution: 70,000, target value: 3e6, maximum injection time: 20 ms) were acquired in the Orbitrap. The 10 most intense ions were sequentially isolated, fragmented by higher energy C-trap dissociation (HCD) and scanned in the Orbitrap mass analyzer (resolution: 35,000, target value: 1e5, maximum injection time: 120 ms, isolation window: 2.6 m/z). Precursor ions with unassigned charge states, as well as with charge states of +1 or higher than +7, were excluded from fragmentation. Precursor ions already selected for fragmentation were dynamically excluded for 25 s. MaxQuant analysis was performed as described above, without setting phosphorylation (STY) as a variable modification. Potential contaminants, reverse hits, hits only identified by site and hits with no unique peptides were excluded from the analysis.

### Atomic force microscopy

Experiments were performed using an AFM setup (CellHesion 200; JPK Instruments) mounted on an inverted microscope (Observer Z1; Carl Zeiss Microscopy). Optical images were recorded with a Plan-Apochromat (25x/0.8NA) water immersion objective. AFM cantilevers with nominal spring constants of 150 mN/m and 350 μm in length (NSC12/tipless/No A1/MikroMasch) were modified with polydimethylsilane (PDMS; Sylgrad 184; Dow Corning) wedges to correct for tilt angle of the cantilever^69^. The spring constant of wedged cantilevers were determined using the thermal noise method^70^. Cells were seeded and grown over night in cover glass bottom dishes (WPI). 2 μM (+)-S-trityl-l-cysteine (STC, Sigma Aldrich) was added 30 min before starting measurements. For each confinement measurement an isolated mitotic cell was identified and DIC imaged before, during and after confinement by the wedged AFM cantilever. Optical images were used to determine cell diameter. The wedged end of the cantilever was lowered onto the glass bottom in the vicinity of the cell. Using a programed procedure, the end was first raised 30 μm at 5 μm/s, held at this height for 20 s while the end of the cantilever was manually positioned above the cell, lowered 22 μm at 1 μm/s to confine the cell. The height of the cantilever was maintained for 60s of confinement, before the cantilever was raised 22 μm at 1 μm/s and the end was repositioned above the glass bottom and finally lowered on to the glass at 5 μm/s. Cantilever height and the force acting on the cantilever were recorded at 100 Hz and the data was processed using JPK analysis software and IGOR (WaveMatrics). Cell diameters were determined using Axiovision (Zeiss) or ImageJ. Cortex tension was determined using a fluid filled model of rounded mitotic cells ^71^.

### Data presentation and statistical analysis

Statistical analysis and visualization of mass spectrometry data MS were performed using the R software environment (version 1.3.1093). GO terms analysis was performed on CUL4B interactors with at least 2-fold enrichment in interaction with WT versus NP1 mutant using the R package ViseaGO^72^ with full human UniProt proteome used as a background. Quantification of images and movies was performed with ImageJ or IGOR (WaveMatrics), which was also used for the analysis of the AFM data. GraphPad Prism was used to generate graphs and perform statistical analysis. For experiments comparing to conditions t-tests were performed, while for multiple conditions 1-ANOVA was used followed by Dunnett’s multiple comparisons test.

## Supplementary tables

Table S1. A list of all phosphorylation sites on CUL4B isolated from a mitotic lysate by immunoprecipitation and TiO2-based enrichment and measured by LC-MS/MS (see Figure 2D).

Table S2. A list of all proteins identified in the triple SILAC interactome of WT CUL4B, NP1 and P50L mutants isolated from a mitotic lysate by immunoprecipitation and measured by LC-MS/MS, with indicated fold change in the interaction between WT and the mutants (see Figures 4B and 6A).

Table S3. GO biological processes of proteins preferentially interacting with phosphorylated CUL4B in comparison to the NP1 mutant (see Figure 6B).

