## Extended Data Figures for "The CRL4B E3 ligase regulates mitosis by recruiting phospho-specific DCAFs"

### Extended Data Figures titles and legends

EDF 1

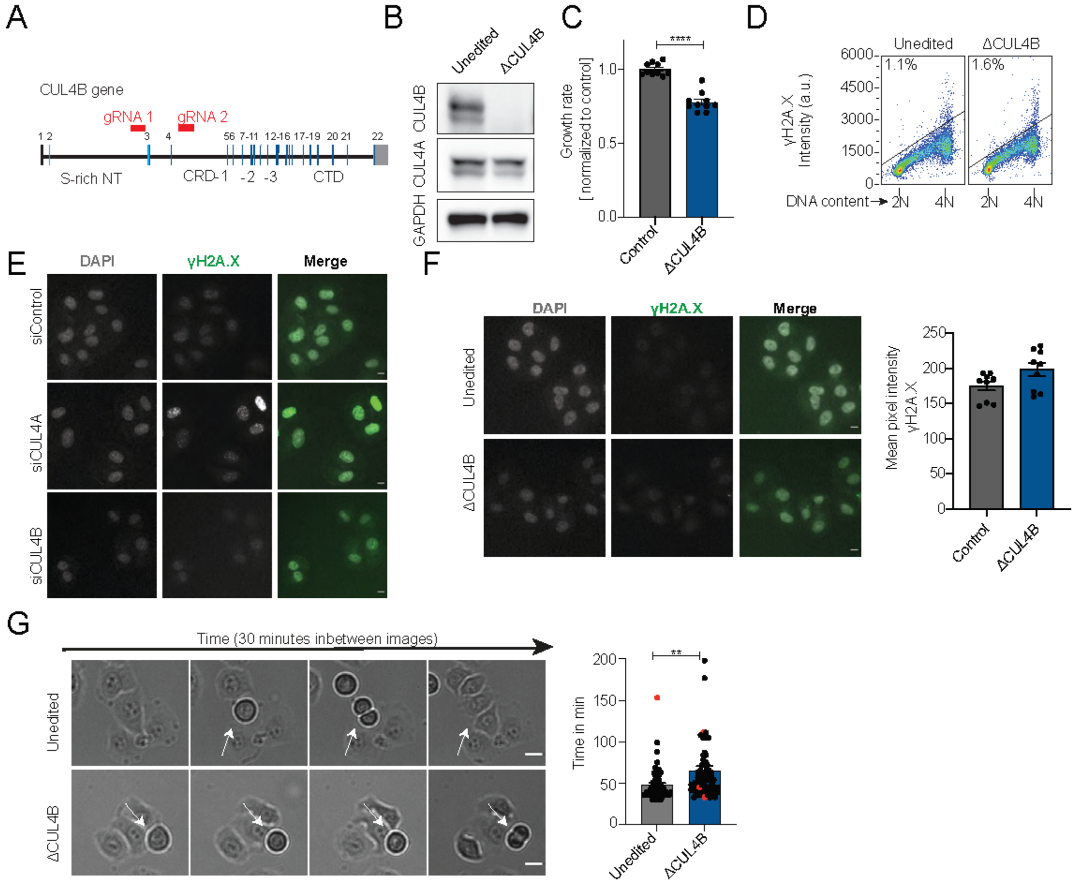


**EDF 1: CUL4B-specific functions in mitosis.** (**A**) Schematic representation of the CUL4B gene with the gRNAs target regions used to delete CUL4B by CRISPR/Cas9 (ΔCUL4B) highlighted by the red bars. The light blue lines depict exons coding for the N-terminal region, while the dark blue ones represent coding exons of the CUL4B core. (**B**) Immunoblotting confirms lack of CUL4B but not CUL4A protein expression in extracts prepared from unedited control or ΔCUL4B HeLa Kyoto cells. GAPDH controls for equal loading. (**C**) The growth rate of ΔCUL4B cells was measured by MTT assays and normalized to unedited controls after 72 h. Mean with SEM of N=3 x 3 technical replicates are depicted. (**D**) Flow cytometry analysis of control cells and ΔCUL4B cells stained γH2A.X and DNA staining with propidium iodide N=3. (**E**) Representative images of DNA damage analysis in RNAi-depleted cells. The same contrast settings were used for representative images of each channel. Scale bar: 10 µm. (**F**) DNA damage levels were analyzed by γH2A.X immunofluorescence in

ΔCUL4B und unedited control cells. The same contrast settings were used for representative images of each channel, and the mean grey value of the γH2A.X signal in the nucleus was quantified. Scale bar: 10 µm. The graph displays mean with SEM of N=3 x 3 technical replicates of at least 50 cells in each experiment. (**G**) Live cell imaging of ΔCUL4B cells. Representative images show intervals of 30 min of the brightfield channel. Quantification of mitotic duration was assessed with brightfield imaging every 3 min counting frames from cell rounding to telophase. Mean with SEM and single values are depicted of N=3 of at least 10 cells in each experiment. Red symbols mark cells with incomplete mitosis during the movie. Scale bar: 10 µm. A t-test was used to assess statistical significance.

EDF 2

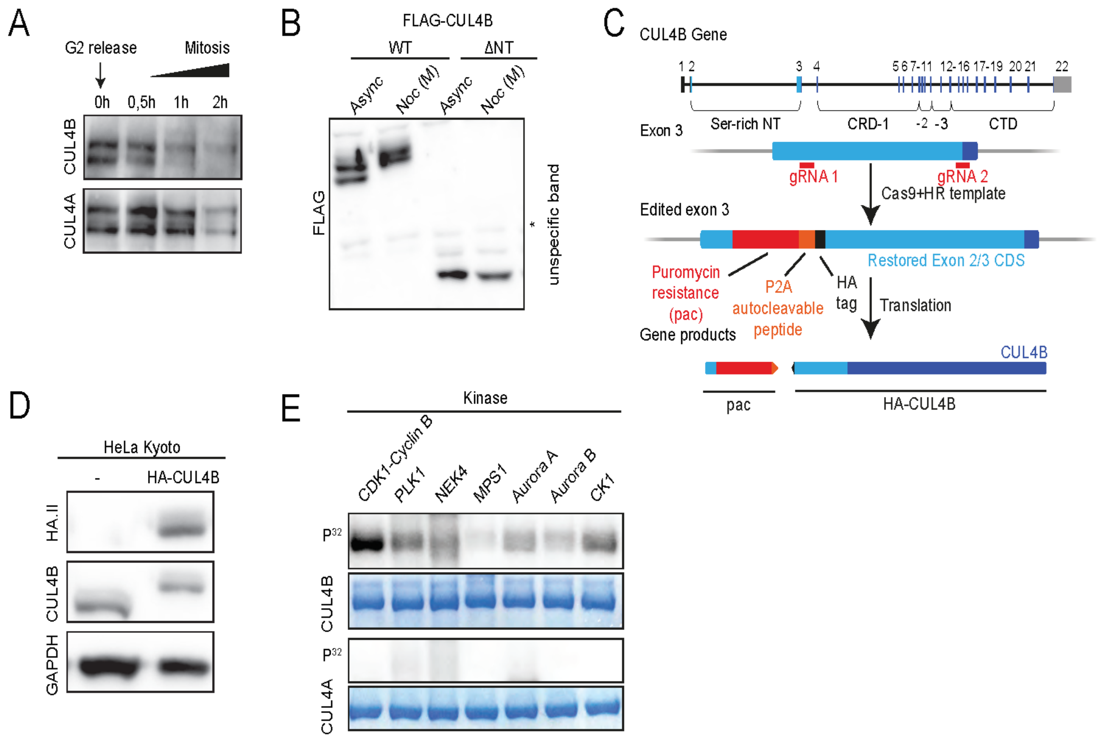


**EDF 2: N-terminal phosphorylation of CUL4B during mitosis.** (**A**) CUL4A and CUL4B were immunoblotted of extracts prepared from cells synchronously entering mitosis after a G2 block/release (RO-3306 for 20 h) (N=2). (**B**) Cells were transfected with the indicated FLAG-CUL4B constructs, and the migration pattern was analyzed. ΔNT lacks the N-terminal region. (**C**) Schematic representation of knock-in strategy of HA-tagged CUL4B. Top: *CUL4B* gene with coding exons. Light blue lines depict exons coding for the N-terminal region, while dark blue lines represent coding exons of the CUL4B core. Middle: representation of Exon 3 before and after editing. The gRNAs targeting regions are marked with red bars and the insert after homologous recombination is depicted. The color assignments for the insert corresponds to the gene product shown at the bottom. Red: puromycin resistance cassette, orange: P2A cleavage peptide, black: HA-tag[73]. (**D**) Immunoblotting of extracts prepared from unedited (-) or edited (HA-CUL4B) HeLa Kyoto cell lines, using HA.II (top panel) or CUL4-CT (middle panel) antibodies. GAPDH controls equal loading (bottom panel). Note that the endogenous CUL4B locus was successfully replaced by HA-tagged CUL4B. (**E**) *In vitro* kinase assay in the presence of $[\gamma\text{-}^{32}P]$ATP using Sf9-purified CUL4A (CRL4A; lower panel) or CUL4B (CRL4B; upper panel) in complex with DDB1

and RBX1 as substrates for a panel of purified kinases. Phosphorylation was monitored by autoradiography ($P^{32}$; upper panels), while equal loading was confirmed by Coomassie blue staining of the SDS-PAGE gel (N=2).

EDF 3

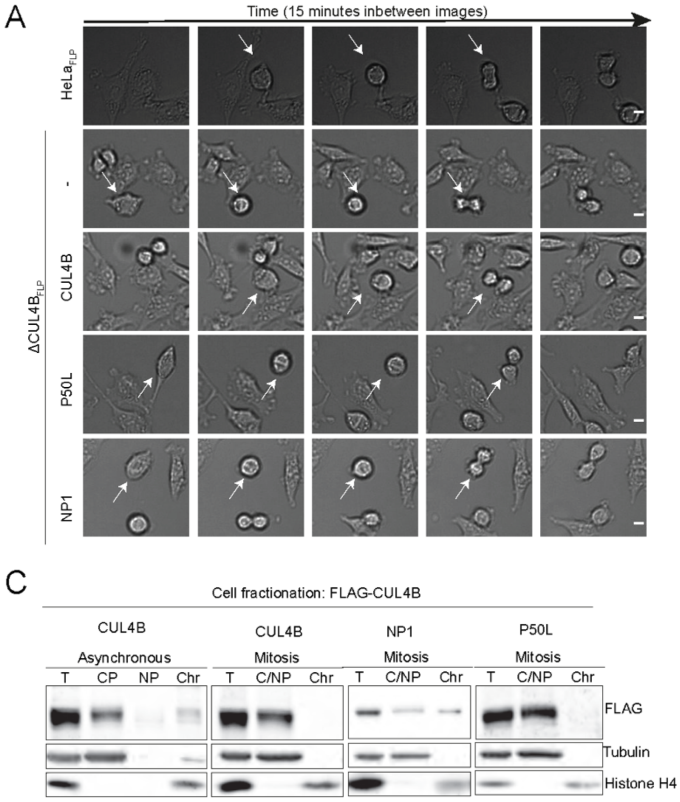


**EDF 3: Mitotic CRL4B phosphorylation promotes progression through mitosis and regulates chromatin association.** (**A**) Brightfield images of life cell imaging of Hela$_{FLP}$ control cells and ΔCUL4B$_{FLP}$ cells expressing either no CUL4B (-), CUL4B, or the NP1 and P50L mutants after doxycycline induction. Representative images of cells (white arrows) progressing through mitosis are shown at 15 minutes intervals. Scale bar: 10 µm (**B**) Cell fractionation experiments of asynchronous and nocodazole-arrested (mitosis) control cells (upper panels, siControl) and DDB1 RNAi-depleted cells (lower panels). Total extracts (T) were fractionated using differential centrifugation, and the distribution of CUL4A, CUL4B and DDB1 was analyzed by immunoblotting. Antibodies against Tubulin and Histone H4 serve as controls for the cytoplasmic and chromatin fractions, respectively. CP: cytoplasm; NP: Nucleoplasm; Chr: chromatin; C/NP: combined cytoplasm and nucleoplasm in mitotic cells. (**C**) Cell fractionation experiments in asynchronous and nocodazole-arrested cells (mitosis) expressing Flag-tagged CUL4B or as indicated the NP1 or P50L mutants (N=3 for

FLAG-CUL4B and P50L, N=1 for NP1). The fractionation and analysis were performed as described in panel B, except that the CUL4B was detected by FLAG-antibodies.

EDF 4

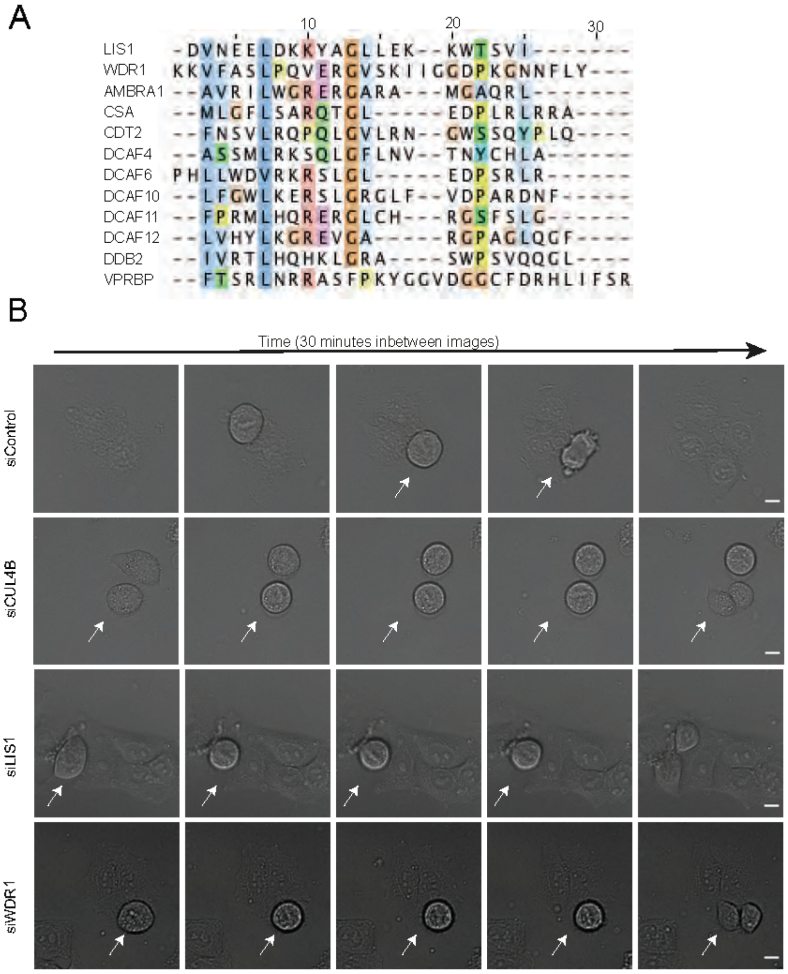


**EDF 4: LIS1 and WDR1 contain a predicted helix-loop-helix motif and may alter mitosis.** (**A**) Alignment of a conserved helix-loop-helix motif of LIS1 and WDR1 with multiple *bonafide* DCAFs. (**B**) Live cell imaging of mitosis in HeLa Kyoto cells, RNAi-depleted as indicated for CUL4B, LIS1 or WDR1, or treated with control si-all star negative oligos (siControl). Brightfield images were taken every 30 min, and mitotic cells marked by the white arrow. Scale bar: 10 µm

EDF 5

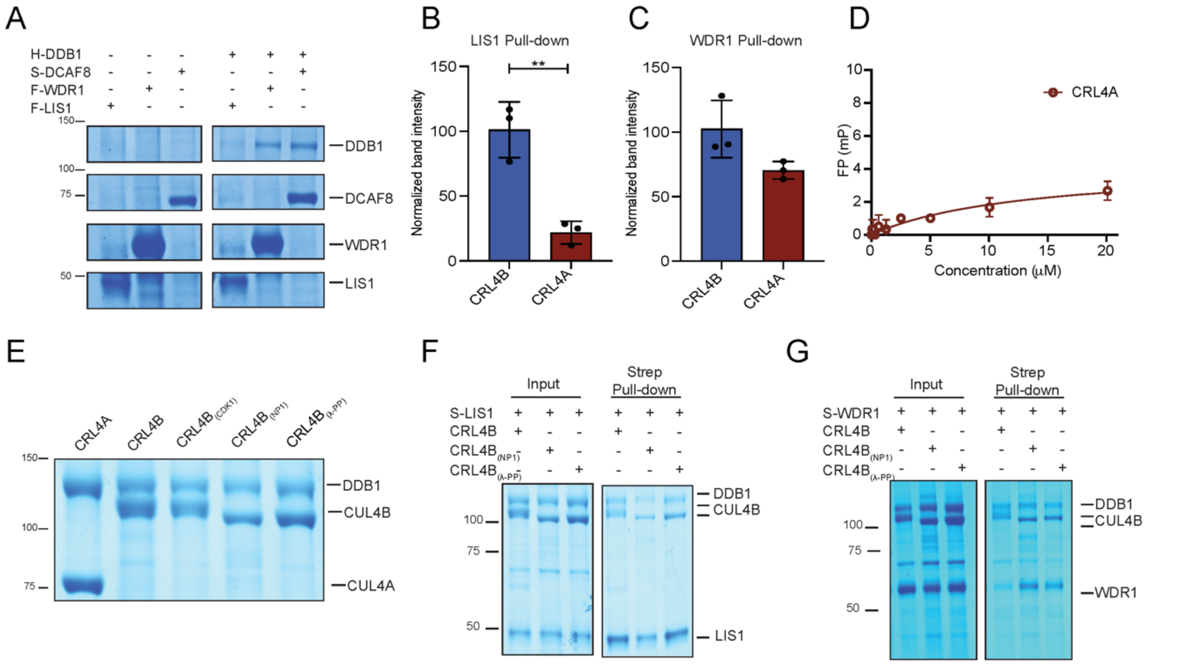


**EDF 5**: **Recruitment of LIS1 and WDR1 into the CRL4B complex is mediated via two different binding sites, one on DDB1 and one on phosphorylated CUL4B.** (**A**) Coomassie-stained SDS-PAGE of *in vitro* pull-down assays of FLAG-tagged LIS1 and WDR1, and His-DDB1, co-expressed in baculoviral *Sf9* cells (N=3). For control, binding of Strep-DCAF8 to His-DDB1 was included (N=2). Note that DDB1 directly interacts with LIS1 and WDR1. (**B**) and (**C**) *In vitro* binding assays (Strep-pull down) of Strep-LIS1 (**B**) and Strep-WDR1 (**C**) with purified CRL4A or CRL4B. Band intensities were quantified and normalized to LIS1 and WDR1, respectively (N=3 for LIS1; N=2 for WDR1). A t-test was used to assess statistical significance. (**D**) Fluorescence polarization (FP) assay (with increased concentrations compared to the experiment shown in Figure 5) quantifying binding of CRL4A to Alexa-labeled LIS1. (**E**) Coomassie-stained SDS-PAGE of Sf9-purified CRL4B and CRL4B treated with CDK1 ($CRL4B_{CDK1}$) or dephosphorylated with λ-PP ($CRL4B_{\lambda\text{-}PP}$). CRL4A and the phosphorylation-defective NP1 mutant ($CRL4B_{NP1}$) served as controls. Note that CUL4B migrates faster after dephosphorylation with λ-PP, implying that a large fraction of CUL4B is phosphorylated when purified from Sf9 cells. (**F**) and (**G**) Coomassie-stained SDS-PAGE of purified proteins (input) and *in vitro* binding assays (Strep-pull down) of Strep-LIS1 (**F**) and Strep-WDR1 (**G**) with purified wild type

CRL4B, the NP1 phospho-mutant ($CRL4B_{NP1}$), or λ-PP treated, dephosphorylated CRL4B ($CRL4B_{\lambda\text{-}PP}$).

EDF 6

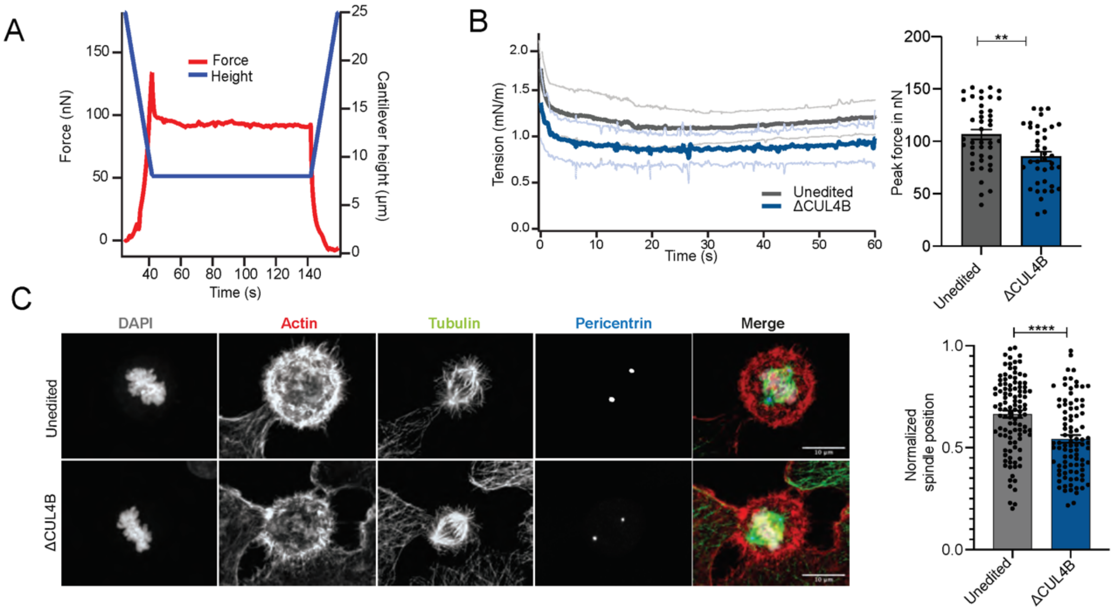


**EDF 6: CRL4B$^{LIS1}$ and CRL4B$^{WDR1}$ complexes regulate cortex tension and spindle positioning during mitosis.** (**A**) Representative mitotic confinement assay. Diagram indicates wedged cantilever end height (µm, blue) and upward force acting on cantilever (nN, red) over time (s) during the confinement phase of the assay. (**B**) Mean cortical tension and the peak force in unedited or ΔCUL4B cells during mitosis. Confinement assay was performed after 30 min of (+)-S-trityl-l-cysteine (STC) treatment. Cortical tension graph (left) shows the mean tension with SD (mN $m^{-1}$) of cells. The peak force (right) (in nN) is given for each cell and the mean with SEM (N=3, including 15 cells). (**C**) Immunofluorescence staining of actin, tubulin and the centrosome component pericentrin was used to visualize spindle positioning in unedited control and ΔCUL4B cells. Representative confocal images show maximum intensity Z-projections of the acquired Z-stacks. The contrast settings are identical between images of a single channel. DNA was visualized with DAPI, and the merge shows the overlay of all channels. Scale bar: 10 µm. Spindle positioning was assessed with ImageJ and the mean with SEM and single values are plotted (N=3, including 50 cells). Statistical analysis was performed with a t-test.
